# CBD resistant *Salmonella* strains are susceptible to Epsilon 34 phage tailspike protein

**DOI:** 10.1101/2022.10.06.511232

**Authors:** Iddrisu Ibrahim, Joseph Atia Ayariga, Junhuan Xu, Ayomide Adebanjo, Michelle Samuel-Foo, Olufemi S. Ajayi

**Author notes:** Correspondence (J.A.A.); (O.S.A.). (I.I.). (J.A.A); (A.A.) (J.X.); (M.S.-F.); (O.S.A.).

## Abstract

The rise of antimicrobial resistance is a global public health crisis that threatens the effective control and prevention of infections. Due to the emergence of pandrug-resistant bacteria, most antibiotics have lost their efficacy. Meanwhile, the development of new antimicrobials has stagnated, which leads to the creation of new and unconventional treatments. Bacteriophages or their components are known to target bacterial cell walls, cell membranes, and lipopolysaccharides (LPS) and hydrolyze them. Bacteriophages being the natural predators of pathogenic bacteria, are inevitably categorized as “human friends”, thus fulfilling the adage that “the enemy of my enemy is my friend”. Leveraging on their lethal capabilities against pathogenic bacteria, researchers are searching for more ways to overcome the current antibiotic resistance challenge. Bacteriophages are considered to be one of the most effective alternative therapies for multidrug resistant bacteria. In this study, we expressed and purified epsilon 34 phage tailspike protein (E34 TSP) from the E34 TSP gene which was previously cloned into a pET30a-LIC vector, then assessed the ability of this bacteriophage protein in the killing of two CBD-resistant strains of *Salmonella* spp. We observed that the combined treatment of CBD-resistant strains of *Salmonella* with CBD and E34 TSP showed poor killing ability whereas the monotreatment with E34 TSP showed considerably higher killing efficiency.

## 1.0 Introduction

Cannabidiol (CBD), a constituent of the *Cannabis sativa* (hemp) plant is a non-psychoactive biochemical extract that plays an essential role in health and physiology. CBD has worldwide applications in medicine and possesses tremendous potential for pharmaceutical relevance, such as anti-microbial, antioxidants, anti-inflammatory, anti-cancer, and anti-convulsant potentials among many others. Several studies have reported the particularly enormous roles CBD plays in antimicrobial (anti-parasitic, antiviral, and antibacterial) infections (**Seltzer et al., 2020**). **Long et al., 2022** described in their studies how CBD was used to inhibit SARS-COV 2 infection in cells and mice by blocking viral gene expression after gaining access to host cells. Their study relied on CBD and its metabolites 7-OH-CBD but did not include Tetrahydrocannabinol (THC). In other works, the anti-viral properties of CBD has been reported, particularly on SARS-COV2 (**Fernandes et al., 2022, Liu et al., 2022, McGrail et al., 2022, Janecki et al., 2022**). **Gildea et al., 2022** examined the antibacterial properties of CBD against two strains of *Salmonella* (*typhimurium and newington*) and demonstrated that CBD was able to disrupt the membrane of the *Salmonella* spp. used in the study. Synergistic studies of polymyxin B and CBD co-therapy demonstrated strong antibacterial activities against *Acinetobacter baumannii* ATCC 19606 and other gram-negative bacteria (*Klebsiella pneumoniae* and *Pseudomonas aeruginosa*) (**Hussein et al., 2022**).

As a non-psychoactive agent, CBD demonstrates its anti-angiogenic, anti-proliferative, and anti-apoptotic properties using several molecular mechanisms, which do not involve signaling its counterpart (cannabinoid) receptors (CB1 and CB 2), and vanilloid receptor 1. CBD’s biological prowess is by stimulation of the endoplasmic reticulum (ER) stress and inhibiting AKT/mTOR signaling. This activates autophagy and stimulates apoptosis. Additionally, CBD promotes the development of reactive oxygen species (ROS), to further invigorate apoptosis. Furthermore, CBD molecular transduction includes upregulating the expression of intracellular adhesion molecule 1 (ICAM-1), as well as tissue inhibition of cell-matrix metalloproteinases-1 (TIMP1). This decreases the expression of inhibitors of DNA binding 1 (ID-1). CBD also activates transient receptors of possible vanilloid type 2 (TRPV2), which promotes the uptake of various cytotoxic agents in cancer cells (**Peyravian et al., 2022, Adams et al., 1940**).

A whopping 115 million human infections and 370,000 deaths per annum are attributed to *Salmonella* infections globally. *Salmonella* is an important foodborne pathogen classified into 2,659 strains based on their surface antigens. *Salmonella typhimurium* is among the most debilitating strains of health uproar (**Qin et al., 2022**). S. *typhimurium* is estimated to have resulted in 16-33 million infection cases and about 500,000-600,000 deaths globally per annum (**Gut et al., 2018**). Though antibiotics play an enormous role in mitigating *Salmonella* infections, their efficacy is becoming futile due to the ability of bacteria to develop stringent resistance to antibiotics. This ontogeny has become a canker to health, and treatment of *Salmonella* infections. This foreclosure of *Salmonella* ontogeny means that novel antibiotic therapies must be fabricated to meet the urgent global health demand (**Rogers et al., 2021, Balasubramanian et al., 2019**).

The natural enemies of bacteria are the bacteriophage. They can attack their target bacteria and are generally host-specific. This specificity attribute of phages allows for the maintenance of the human host’s microbiota if used as therapy. However, phages are self-limiting, easily cleared from the body, elicits strong immune response, thus making it problematic as a therapeutic agent. Additionally, these phages have narrow host range, thus compounding their use as antibacterial agents against a broad spectrum of bacterial infections. In phage therapy, it is crucial that the preparation of the stocks is free of bacterial toxins such as the bacteria LPS and even the host bacteria as a whole since these by themselves are agents of disease. The elimination of bacterial toxins and bacteria during the preparation steps thus presents a technical challenge and increases the production cost (**Rahman et al., 2021, Borysowski et al., 2006, Fischetti et al., 2005, López et al., 2004**). To overcome these huddles, a much cheaper and safer approach could be the utilization of the phage’s hydrolases which the phage uses as its host attacking machinery.

The E34 phage belongs to the P22-like phages which have a uniquely short tailspike architecture, thus the name podovirus. Their tailspike protein is characterized by the presence of a globular head binding domain, a parallel beta-helix domain, and a beta-prism domain (**Kim, 2006**). Although the structures that are involved in viral adhesion and infection vary, a broad generalization of their properties can be made due to the presence of the β-helix domain. This unique domain is found in most of the viral and fungal structures that are commonly used in the development of human infectious diseases. Some of these include those of *Chlamydia, Helicobacteria, Borrelia*, and *Rickettsia*. The parallel β-helix domains of other bacterial and fungal proteins are also known to be found in various surface and bacterial proteins.

Usually, parallel β-helix folds are not found in higher eukaryotes. However, these structures are known to exhibit high melting point (Tm), stability, and resistance to detergents at room temperature. Their role in the development of exterior virion structures that can endure harsh environmental conditions is also evidenced by their properties.

Most β-sheet proteins have been studied. The first known example of this is the pectate lysases from the Erwinia species, which are known to have a unique structure that allows them to infect plant cells. They are characterized by a long-arm elongated or solenoid-shaped structure. Most of the P22-like protein families, such as SF6, E15, and P22, have a unique structure that allows them to recognize and attach to host receptors. This structure also allows them to develop an LPS cleaving mechanism. The economic and sanitary burden caused by antimicrobial resistance (AMR) is immense.

An in-depth analysis conducted by a group of researchers revealed that the number of deaths caused by bacterial infections could be as high as 4.95 million annually (**O’Neill, 2016, Huttner et al., 2013, Laxminarayan et al., 2013**). The researchers used a statistical model to analyze 471 million records from 16 different countries. They found that the mortality rate from bacterial infections could be as high as 1.27 million annually (**Smith and Coast, 2013, Murray et al., 2022**). One of the most common causes of deaths associated with antimicrobial resistance is intra-abdominal infection. This contributes to the economic impact of the issue, as it costs the United States around USD 20 billion annually.

This study employs the tailspike proteins (TSP) of epsilon 34 (E34) phage that was previously expressed and purified by **Ayariga et al., 2022** in combination with CBD to understand its interactions with CBD-resistant *Salmonella typhimurium* and *Salmonella newington*. In this work, we performed plate assays, fluorescence microscopy, and growth kinetic studies to determine the antibacterial activity of CBD extracted from *C. sativa* in combination with E34 TSP on resistant strains of *S. typhimurium* and *S. newington*. Additionally, we conducted comparative kinetic studies of the two *Salmonella* strains in the presence of CBD, E34 TSP, and their combinations. We hypothesized that Phage protein-CBD combination exhibits antibacterial activity against *S. typhimurium* and *S. newington*. The results of these studies suggest that phage-CBD does exhibit potent antibacterial activity against *Salmonella* typhimurium and not *Salmonella newington*. This rather unpredicted outcome warrants further encouraging research and development of CBD/phage as a potential antibacterial agent.

## 2.0 Materials and Methods

### 2.1 Media, chemicals, bacterial strains, and other reagents

The transformants were grown on agar medium, which was premixed with 50 mg/mL of kanamycin antibiotic. A final concentration of 1 mM isopropyl-β -D-thiogalactopyranoside (Sigma) was used for induction. All media, enzymes and oligonucleotides for PCR reaction, transformation and induction were purchased from New England Biolabs. Competent cells BL21/DE3 and Novablue cells were purchased from Novagen, P22 host cells, *Salmonella typhimurium* was from our laboratory collection, *Salmonella newington* was a kind gift from Andrew Wright (Tufts) and Sherwood Casjens (Utah). pET30a-LIC vector was purchased from Novagen, urea, sodium chloride, ammonium sulfate, and all other chemicals used in this research were of HPLC grade. Two strains of *Salmonella* were used in this study. They are lab coded as BV4012 and BV7004 representing *Salmonella typhimurium* LT2 strain MS1868 (a kind gift from Dr. Anthony R. Poteete, University of Massachusetts) and *Salmonella newington* as *S. newington* (also known as *S. enterica* serovar Anatum var 15+ strain UC1698, a kind gift from Dr. Sherwood R. Casjens, University of Utah) respectively. The *S. typhimurium* and the *S. newington* were subjected to prolonged CBD doses to cause resistance development.

The resistant colonies were isolated by plating on LB agar spiked with CBD. We have previously shown that these strains were susceptible to CBD at micromolar concentrations (**Gildea et al., 2022**). These resistant strains were treated to high doses of CBD, E34 TSP, or a combination of both.

### 2.2 E34 TSP expression and purification

The initial cloning of the E34 TSP gene into Pet30a-LIC has been described elsewhere (**Ayariga et al., 2022**). Verification of a cloned insert was obtained by PCR analysis via 0.8 % agarose gel electrophoresis using gene-specific primers.

BL21/DE3 cells containing the pET30a-LIC with the E34 TSP insert were streaked on Luria agar plates containing kanamycin. Streaked plates were incubated at 37 °C in Precision Economy incubator overnight. This vector contains a 6HIS region that can be targeted for HIS-tag purification. Single colonies were selected and grown in LB broth primed with Kanamycin in a MaxQ 4450 incubator (Thermo Scientific) fitted with a shaker running at 121 rpm. At mid-log, IPTG (Sigma) was added to a final concentration of 1mM IPTG to induce cells. An uninduced control sample served as control for subsequent protein analysis. After 6 h, the bacteria were pelleted at 10,000 rpm in an Avanti J XP centrifuge fitted with JA14 rotor chilled at 4 °C. Pelleted cells were then re-suspended in lysis buffer consisting of 50 mM Tris at pH7.4, 5 mM MgCl_2_, 0.1 mg/mL lysozyme, 0.1 mg/mL DNase, 0.05 mg/mL RNASE, 0.2 mg/mL DTT and subjected to three cycles of freeze-thaw-freeze. Samples were then centrifuged at 17,000 rpm for 30 minutes and the supernatant decanted into 50 mL tubes and stored at −20 °C as the E34 TSP lysate. The fractionation of E34 TSP was then carried out using FPLC (GE/Amersham Biosciences-AKTA) connected to a desktop computer Pentium 4 running UNICORN software; fractions were pooled and enriched to the desired concentrations using Amicon concentrators (MilliporeSigma). Purified samples were then run on 10% SDS PAGE to determine the induction and the purity of the protein.

### 2.3 CBD stock preparation and serial dilutions

The CBD hemp variety ‘Suver Haze’ CBD stock was obtained from Sustainable CBD LLC., and the extraction process has been published elsewhere (**Gildea et al., 2022**). The stock was diluted with EtOH and vortexed to a final concentration of 50 mg/mL CBD and 4% EtOH. This was further diluted serially with 4% EtOH to produce our working concentrations of 250 μg/mL, 125 μg/mL, 62.5 μg/mL, 31.25 μg/mL, 15.62, and 7.81 μg/mL of CBD.

### 2.4 Creating CBD resistant strains

To investigate the antimicrobial activity mechanism of CBD, we generated CBD resistance following a procedure published by **Wassmann et al., 2022**. *S. typhimurium* strain BV4012 and *S. newington* strain BV7004 were subjected to CBD resistance development by growing them for an extended time (7 days) in media supplemented with CBD. Initially, low doses of CBD (1-5 μg/mL), followed by selecting resistant colonies and growing them in CBD concentrations of 10-50 μg/mL. Finally, resistant colonies from these were plated on LB agar supplement with 50 μg/mL of CBD. Colonies formed were then grown again in LB broth to reach log phase. Resistant strains were then pelleted and resuspended in 1X PBS, and 50% glycol added. Finally, resistant bacterial samples were aliquoted into microcentrifuge tubes and stored at −80 °C. Subsequently, CBD resistant strains of *S. typhimurium* and *S. newington* were restreaked on LB agar.

### 2.5 Assessments of CBD, E34 TSP, and their combination treatment of *Salmonella* spp

Serial dilutions of CBD stock solutions were carried out to obtain the final concentrations of 250, 125, 62.5, 31.25, 15.62, and 7.81 μg/mL. Two *Salmonella* strains BV4012 (*S. typhimurium*) and BV7004 (*S. newington*) were grown in LB broth to logarithmic growth phase and diluted to OD_600_s of approximately 0.08 and 0.25. The CFUs were determined via plating on LB agar to consist of concentration of approximately of 1×10^4^ and 1×10^8^ CFU/mL. Then, aliquots of 100 μL of the bacterial cells were seeded in a 96-well microtiter plate (Fisherbrand™, Fisher Scientific, Fair Lawn, NJ, USA) at a density of 1 × 10^4^ per mL to mimic early log growth phase of the bacteria, or at a density of 1 × 10^8^ per mL to mimic log phase. Then, 50 μL of each solution of the protein, or the CBD or the combinations was added to the bacterial cells. The experimental groups were treated with 250, 125, 62.5, 31.25, 15.62, and 7.81 μg/mL concentrations of CBD, or 44.5, 22.25, 11.12, 5.56, 2.78, 1.39 μg/mL of E34 TSP, or the combinations of CBD and E34 TSP in varying concentrations. The controls consisted of dH_2_O group (negative control) and 2% SDS group (positive control). The samples were incubated at 37 °C for set time points. The growth kinetics of bacteria were determined by reading their optical densities at wavelength of 600 nm using SpectraMax plate reader ((Molecular Devices SpectraMax^®^ ABS Plus) (Molecular Devices LLC., San Jose, CA, USA)). The experiments were carried out in biological triplicates.

### 2.6 Membrane integrity assessment via Propidium Iodide and SYTO-9 staining

To assess the capacity of CBD, E34 TSP and CBD-E34 TSP combination to disrupt the cytoplasmic membrane integrity, the membrane impermeable fluorescent DNA intercalating dye Propidium iodide (PI) and SYTO-9 were employed which is a quick and accurate method to determine the proportion of dead and live cells in cell cultures. The fluorescence produced by the propidium iodide (PI) when it binds to DNA is used to identify dead cells (**Zhou, 2011**). However, live cells with intact membranes present a barrier to PI, thus only cells with disrupted membranes can be stained with PI. Hence, this is used in assessing membrane integrity and cell death since dead cells will allow the penetration of PI into their cytoplasm. Fluorescence microscopy was utilized to observe the effects of CBD, or E34 TSP, or the combination of CBD and E34 TSP treatment on *Salmonella* cells. CBD resistant strains of *Salmonella* - strains *S. typhimurium* and *S. newington* - were grown to mid-log phase, diluted to cell concentration of 1 × 10^8^ and then treated with CBD, E34 TSP, or the combination of CBD and E34 TSP. Two controls, dH_2_O and 2% SDS served as the negative and positive controls respectively. Samples were then stained with 1 X SYTO-9 and 40 μg/mL of propidium iodine and left at room temperature for 25 min, covered with aluminum foil. Samples were then observed using an EVOS FLC microscope (Life Technologies Corporation, Carlsbad, CA, USA).

### 2.7 Assessment of membrane lysis via genomic DNA migration

Bacteria cell membrane plays several crucial functions; however the most fundamental function is to act as a storage sac for all the cytoplasmic content. Thus, if lysed, the cytoplasmic content which consists of the bacterial proteins and nucleic acid will leak. Taking advantage of this important role of cytoplasmic membrane, we hypothesized that the genomic materials of lysed cells will migrate in agarose gel matrix, whereas bacteria with intact cell membranes will retain their genomic material and thus block the nucleic acid from migrating through the agarose gel’s matrix when subjected to electrophoresis. To investigate the ability of E34 TSP, CBD or CBD-E34 TSP treatment to cause lysis of bacteria membrane, we pelleted 500 μL of *S. typhimurium* that was previously growing at mid-log phase and resuspended the bacteria in 1X PBS. 100 μL of the bacteria sample was aliquoted into centrifuge tubes and 100 μL of 44.5 μg/mL E34 TSP, or 22.25 μg/mL of E34 TSP, or 250 μg/mL of CBD, or 125 μg/mL of CBD were added to the bacteria samples and incubated for 1 h. Subsequently, 30 μL of each treatment sample was loaded into a 0.8% agarose gel and run for 75 minutes at 90 volts. Gel was then stained with Ethidium bromide and DNA bands visualized using ChemiDoc XRS imager.

### 2.8 Bacterial glutamate dehydrogenase activity Assay

The enzymatic activity of *Salmonella* spp. was assayed to assess the effect of CBD, E34 TSP or CBD-E34 TSP combination treatment on the bacterial cells in culture. In brief, *Salmonella* cells growing at mid-log phase were diluted to an OD_600_ of 0.2. Then, 100 μL of the bacterial sample was placed into each well of a 96-microplate and treated to varying concentrations of CBD, E34 TSP or CBD-E34 TSP. Then 50 μL of the non-toxic resazurin was added to each well. The conversion of resazurin to resorufin produced a shape color change. The color change was monitored using SpectraMax plate reader (Molecular Devices SpectraMax^®^ ABS Plus) (Molecular Devices LLC., San Jose, CA, USA) and absorbance readings recorded at 590 nm (**Alonso et al., 2017**).

### 2.9 Statistical analysis

Two independent bacterial colonies were used for these experiments, and the experiments were carried out in triplicates, and results are presented as means ± SEM. P-values of 0.05 were considered significant using student t-test. All statistical analyses were performed, and graphs were plotted on Microsoft Excel (Microsoft 2010), microscopic images were processed using ImageJ (an opensource NIH software).

## 3.0 Results

In our previous studies, we demonstrated the antimicrobial effect of CBD against the Gram-negative bacterium *S. typhimurium* **(Gildea et al., 2022**), and in a separate publication, we also revealed that CBD synergizes with ampicillin and polymyxin B in killing *S. typhimurium* (**Gildea et al., 2022b**). E34 phage protected Vero cells from *Salmonella* infection (**Ayariga et al., 2021**). In this work, E34 phage’s LPS hydrolase, which is the E34 TSP, was expressed from a previously published clone (**Ayariga et al., 2021**), and used in combination with CBD to investigate the killing abilities of the two antimicrobial agents against two bacterial strains of *Salmonella*. The proceeding sections provides data obtained from multiple biological assays carried out to determine the antimicrobial activities of CBD and E34 TSP against CBD-resistant strains of *S. typhimurium* and *S. newington*.

### 3.1 Verification of E34 TSP gene insect in PET30 a-LIC vector and SDS PAGE analysis of expressed E34 TSP

As shown in **Figure 2A, lane 8**, our PCR product produced the 1.818 kbp size insert which is the size of E34 TSP gene. In **Figure 2B**, the expressed E34 TSP migrated to a size consistent with its trimeric molecular weight size (see lane induced lysate and fractionated E34 TSP 1). The purification process carried out ensured the removal of over 90% contaminating proteins as shown in the fractionated E34 TSP1. Induction of E34 TSP was achieved as a thicker band could be visibly seen in the induced lane compared to the uninduced lane where the E34 TSP band could barely be observed.

**Figure 1.**
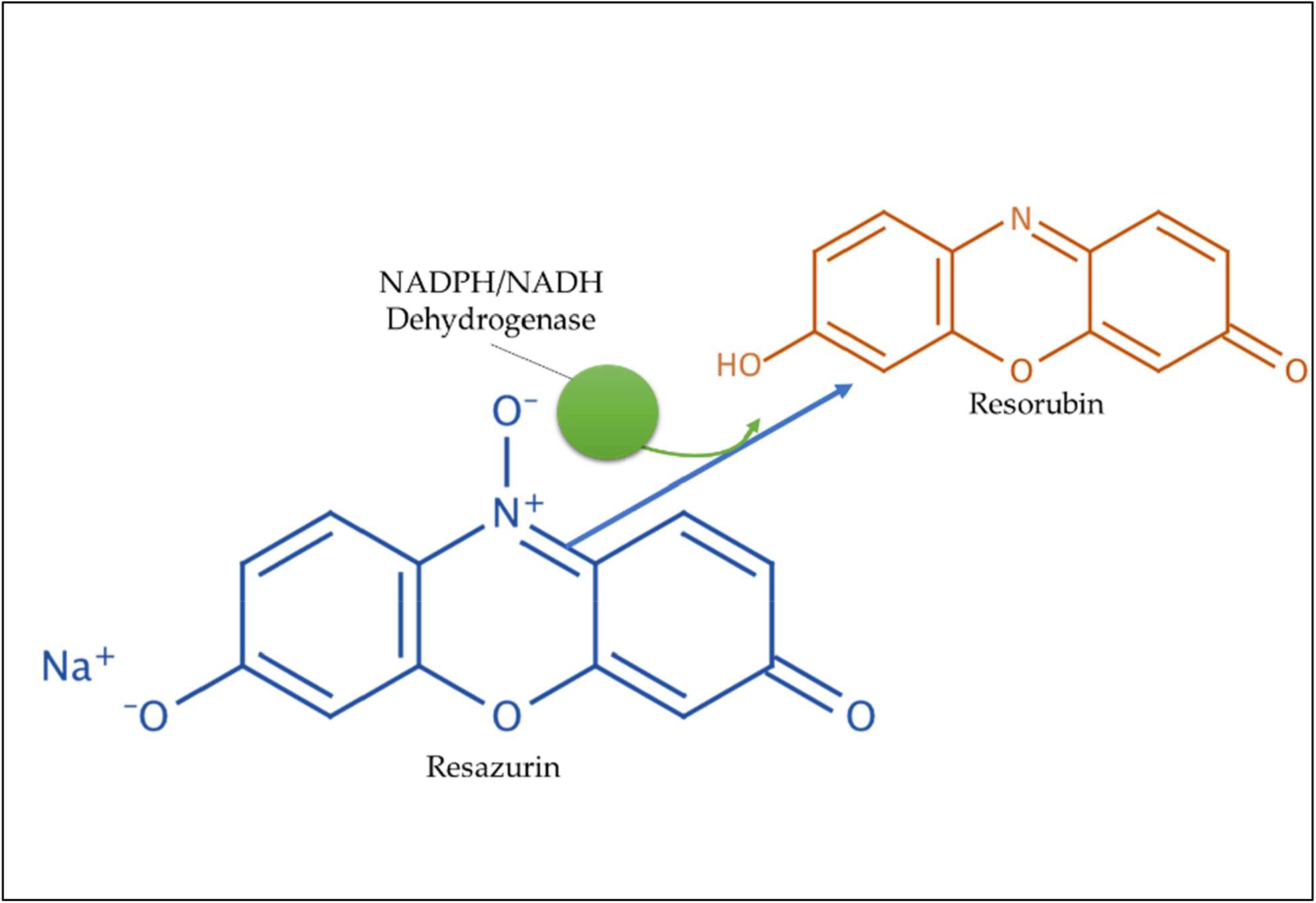
Diagrammatical illustration of the bacterial dehydrogenase activity in converting resazurin to resorubin used as a measure of bacterial metabolism.

**Figure 2.**
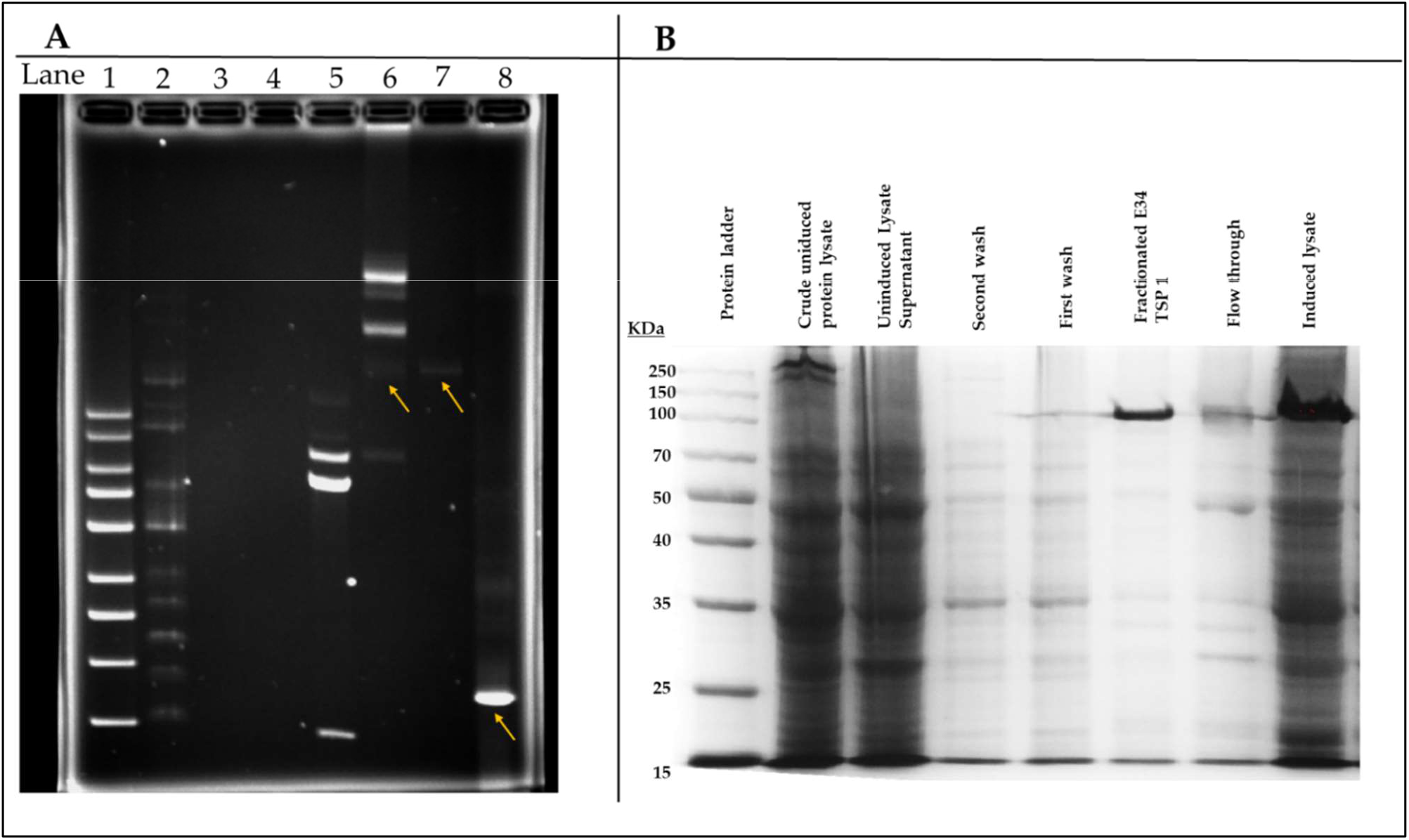
pET30a-LIC-E34 TSP verification and E34 TSP protein purification analysis. **Figure 2A**. Analysis of pET30a-E34 TSP clone in a 0.8% agarose gel. **Figure 2B**. Analysis of E34 TSP lysate and fraction using 10% SDS PAGE. Lane 1 = 1 kb DNA ladder, from Millipore Sigma. Lane 2 = NEB supercoil ladder; New England BioLabs. Lane 3 = miniprep for pET30a-E34 DNA in non-transformed cells. Lane 4 = miniprep for pET30a-E34 DNA in unsuccessfully transformed cells. Lane 5 = miniprep for pET30a-E34 DNA in successfully transformed cells; Nde1 digested pET30a-E34 DNA. Lane 6 = miniprep for pET30a-E34 DNA in successfully transformed cells; undigested pET30a-E34 DNA. Lane 7 = E34 WT DNA. Lane 8 = PCR product from pET30a-E34. **Figure 2B**. SDS-PAGE analysis of E34 TSP protein induced by IPTG, samples separated by 10 % polyacrylamide gel, and visualized via Coomassie brilliant blue R-250. Induction of the E34 TSP was confirmed via the SDS PAGE. Comparing the induced lane and uninduced lane, it is observed that the band representing the E34 TSP is induced and is far denser than what is observed in the non-induced lanes.

**Figure 3.**
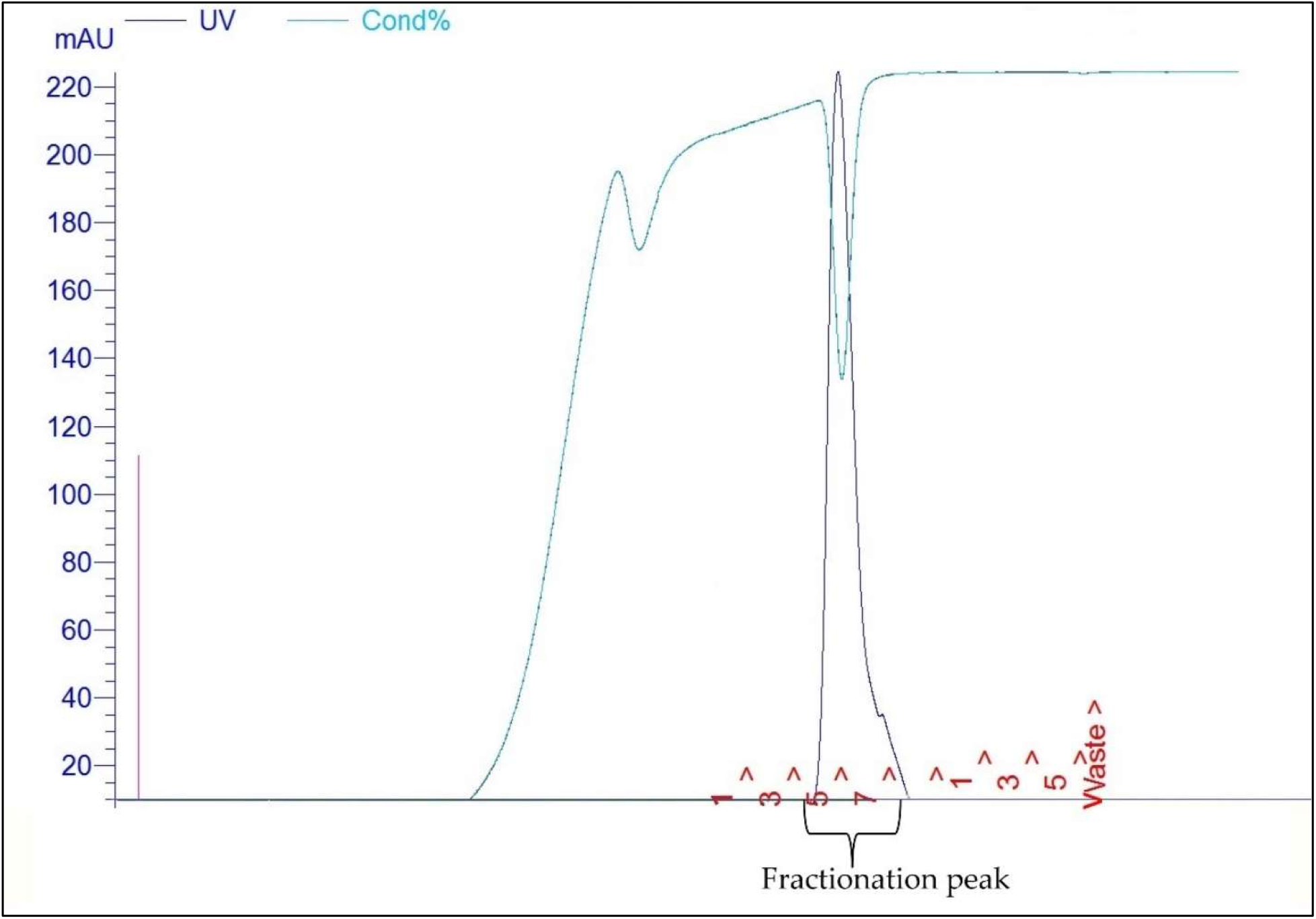
FPLC purification of E34 TSP. Fast protein liquid chromatography (FPLC) chromatograms of E34 TSP: flow rate 1.00 mL/min). Protein fractionations using the FPLC system (Bioscience Amersham), fractionation process was carried out at room temperature. Collected fractions were pooled together, fractions represented those that fell under the fractionation peak corresponding to our protein. The pooled samples were further analyzed via SDS-PAGE as shown in **Figure 2B. As shown in Figure 3**, FPLC fractions of E34 TSP using Co-NTA chromatography was visualized using Coomassie stained SDS-PAGE. Each lane was loaded with equal volume of samples (20 μL each) except lanes 1 which received 10 μL. Lane 1= molecular ladder Colored Prestained (CPPS), Lane 2 = crude uninduced lysate of E34 TSP sample. Lane 3 = Supernatant of the uninduced lysate of E34 TSP. Lane 4 = Second wash. Lane 5 = First wash. Lane 6 = Fractionated E34 TSP Sample. Lane 7 = Flow-through sample. Lane 8 = lysate of prolonged induction of E34 TSP sample. 5 mL Cobalt-NTA FPLC columns (Co-NTA) was used in the FPLC fractionation of E34 TSP samples at pressure of 0.27 MPa, flow rate of 1 mL/minute. Samples that fell under the fractionation peak curve were pooled together and concentrated using Amicon Ultra concentrators (Millipore). Samples of both pooled and concentrates were loaded in to SDS gel to analyze for purity.

### 3.2 Effect of E34 TSP and CBD treatments on *S. typhimurium* and *S. Newington* growth at early and late log phases

At log phase, most bacterial disease symptoms begin to surface, and it is the growth phase which shows most dramatic changes in both bacteria number and disease severity. To assess the effect of E34 TSP, CBD and CBD-E34 TSP combination treatments on *S. typhimurium* and *S. newington* growth at early and late log phases, cells growing at early or late log OD_600_’**s** were subjected to E34 TSP, CBD or CBD-E34 TSP combination treatments as depicted in **Figure 4**.

**Figures 4A-B.**
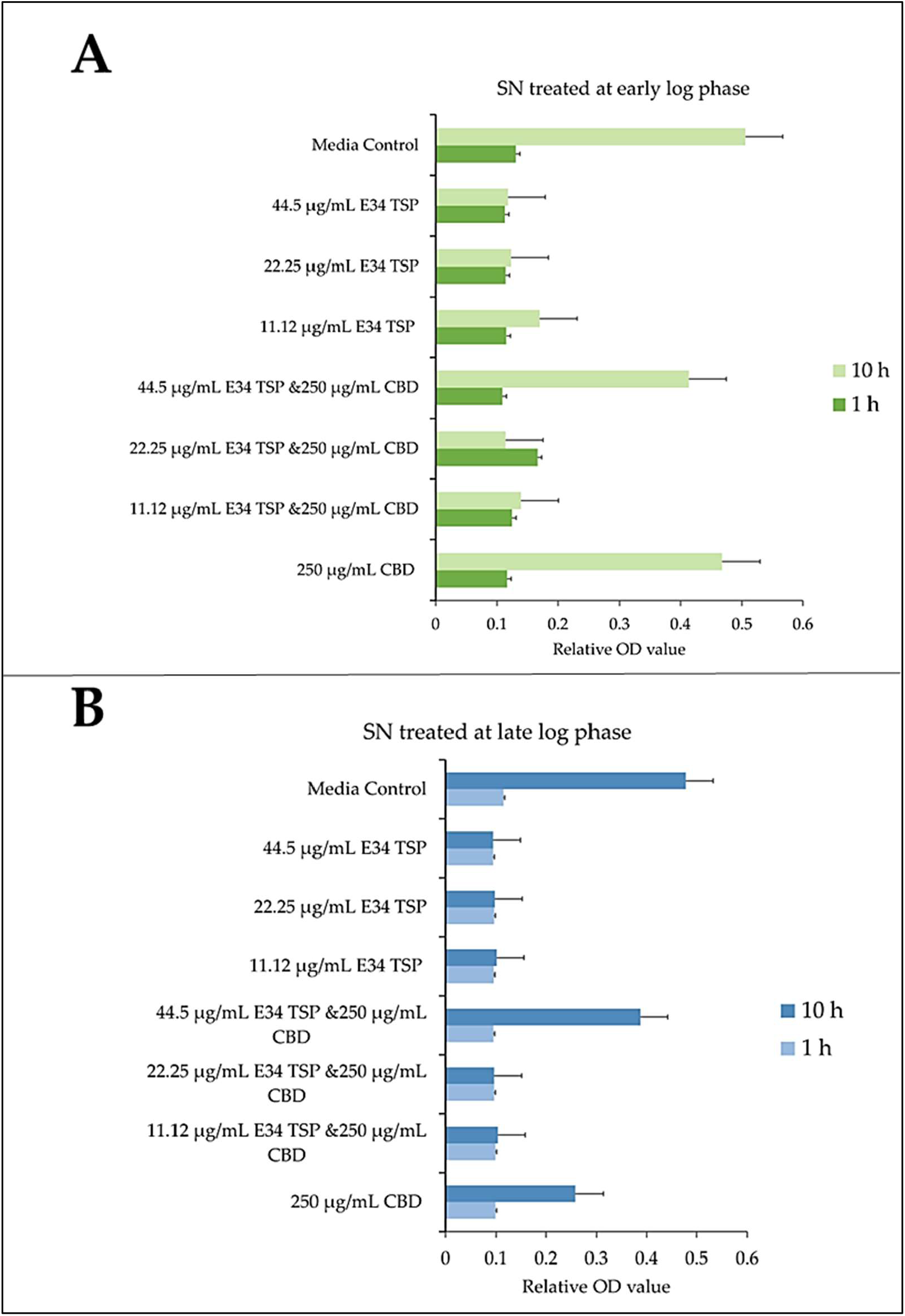
Effect of CBD/E34 TSP treatment of *S. newington* in both early and late log phase of the bacteria species. **(A)** CBD and E34 TSP treatment to *S. newington* (SN) at early log phase. **(B)** CBD and E34 TSP treatment to *S. newington* (SN) at late log phase. Data shown are from three independent experiments expressed as means ± standard deviations.

As shown in **Figures 4A-B**, treatment of *S. newington* with CBD, E34 TSP and the combination of the two produced interesting findings. We observed that the relative OD_600_ of E34 TSP treated samples were generally lower than the combined treatment for the early log-phase at the 10 h time point of culture. The data revealed a sharp decrease in bacterial relative OD_600_s in both E34 TSP treated samples than the combination treatment. The exception however was observed in the 22.25 μg/mL E34 TSP and the 250 μg/mL CBD combination treatment which depicted a drastic decrease in relative OD_600_. The 250 μg/mL CBD treatment only produced relative OD_600_ comparable to the media control. In the 44.50 μg/mL E34 TSP and the 250 μg/mL CBD combination treatment, we observed approximately triple jump in relative OD_600_ as compared to the 44.5 μg/mL E34 TSP monotreatment of *S. newington* **Figure 4A**. Similar trends were observed in the late log phase treatment of *S. newington* to E34 TSP and CBD as depicted in **Figure 4B**.

In **Figures 4C-D**, treating *S. typhimurium* (ST) with E34 TSP showed a relative lower inhibition characteristic after 10 h than the E34 TSP and CBD combination treatments. At higher concentrations of E34 TSP alone, there was a drastic reduction in relative OD_600_, however its combination with CBD performed poorly against the bacteria at the early log phase. Yet again, when E34 TSP was combined with CBD at 44.5 μg/mL and 250 μg/mL respectively, the treatment performed poorly against the bacteria. Treatment of the bacteria with CBD alone did not reduce the relative OD_600_ in the early and late growth phases of the bacteria.

**Figure 4 C-D.**
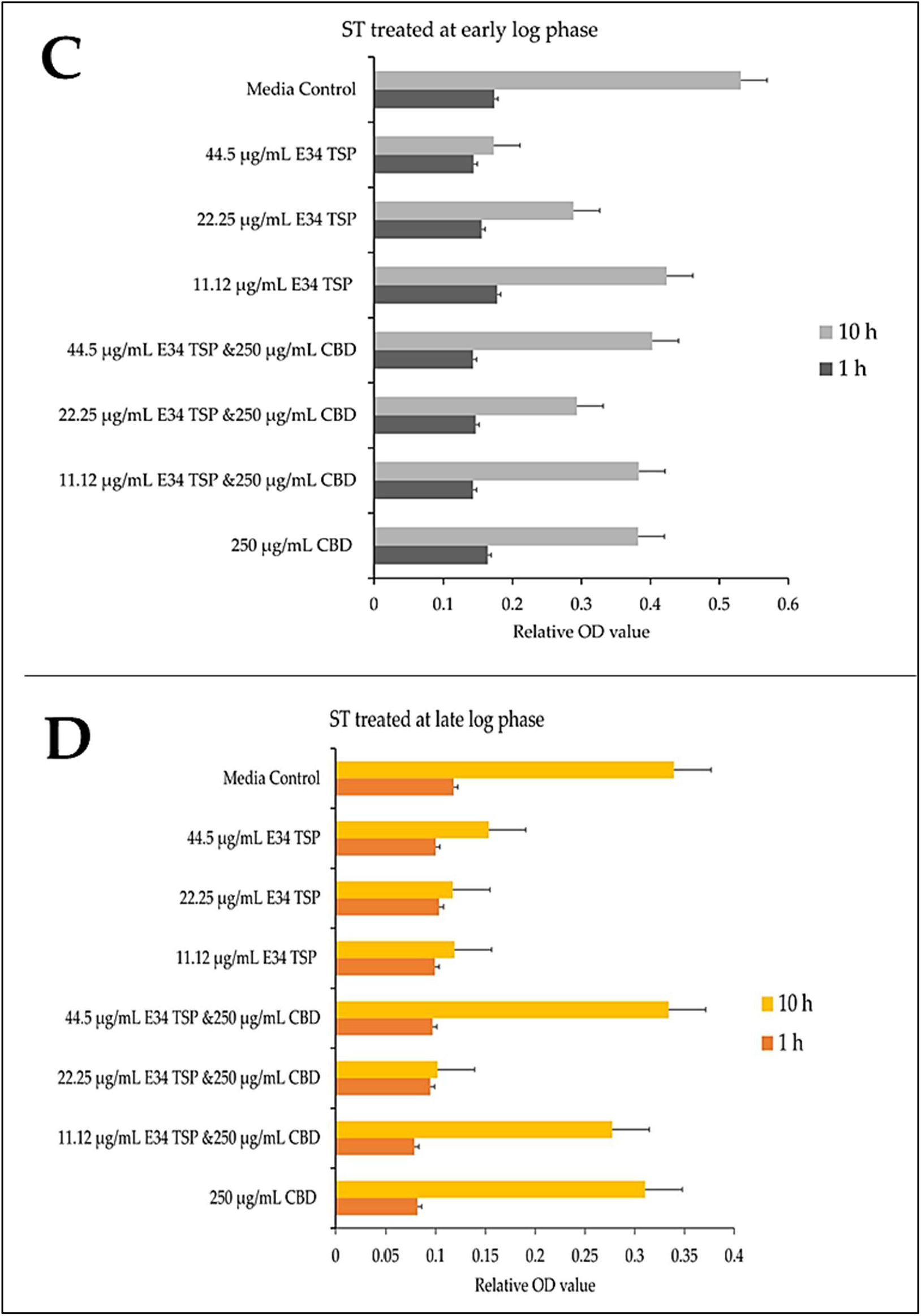
Effect of CBD and E34 TSP treatment of *S. typhimurium* in both early and late log phase of the bacteria species. **(C)** CBD and E34 TSP treatment to *S. typhimurium* (ST) at early log phase. **(D)** CBD and E34 TSP treatment to *S. typhimurium* (ST) at late log phase. Data shown are from three independent experiments expressed as means ± standard deviations.

### 3.3 Immunofluorescent analysis of E34 TSP, CBD, and CBD-E34 TSP combination treatment on *S. typhimurium* and *S. newington*

To investigate the effect of E34 TSP, CBD, and CBD-E34 combination on *S. typhimurium* and *S. newington* viability through the disruption of the bacterial cytoplasmic membrane integrity, the membrane impermeable fluorescent DNA intercalating dye Propidium iodide (PI) and the membrane permeable green, fluorescent dye, SYTO-9 was employed. In brief, CBD resistant strains *S. typhimurium* and *S. newington* were grown to mid-log phase, diluted to cell concentration of 1 × 10^8^ and then treated with CBD, E34 TSP, or the combination of CBD and E34 TSP. Two controls, dH_2_O and 2% SDS served as the negative and positive controls respectively. **Figures 5-9** illustrates the effects of treatment on the viabilities of *S. typhimurium* and *S. newington*.

**Figure 5.**
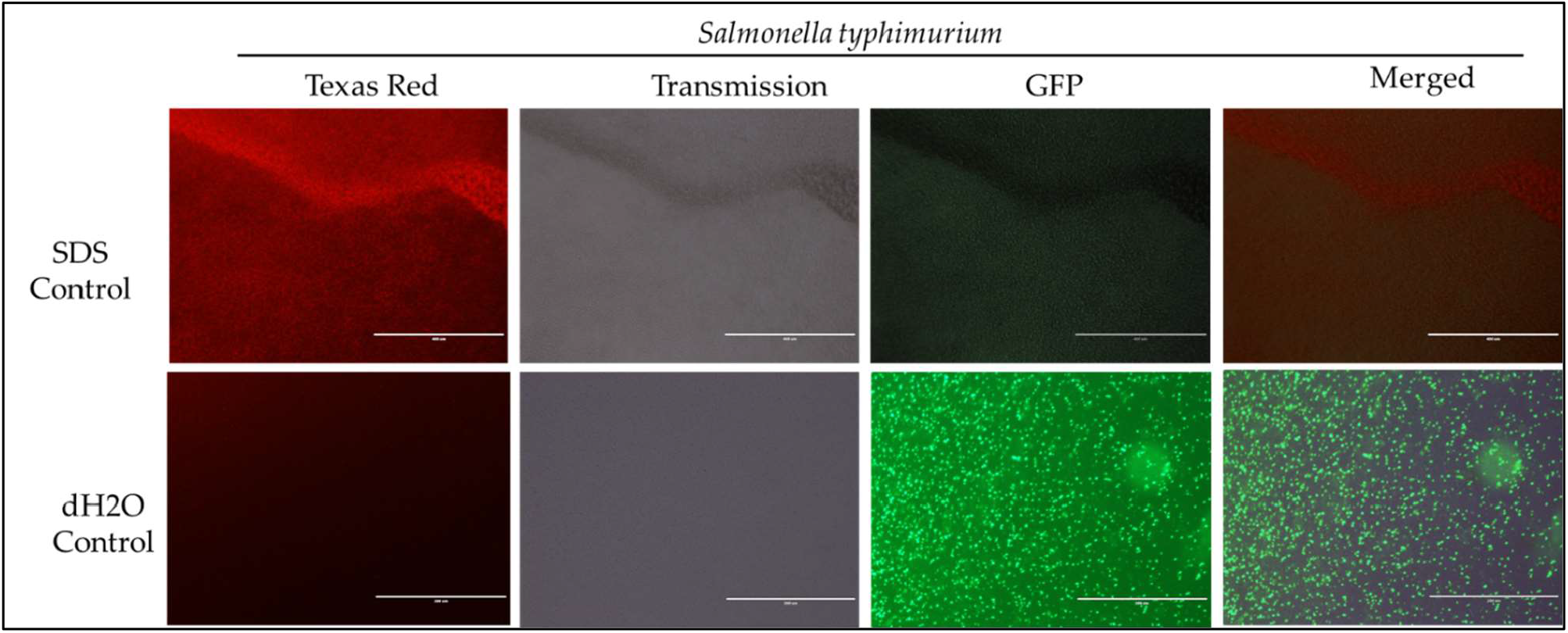
Immunofluorescence analysis of *S. typhimurium* (ST) treated to 2% SDS as a positive control, while dH_2_O as a negative control. Scale bar = 100 μm. The immunofluorescence studies reveal that 2% SDS had killed *S. typhimurium* (as shown in complete red fluorescence), whereas the dH_2_O revealed live bacteria growing in the sample as indicated by the green fluorescence.

**Figure 6.**
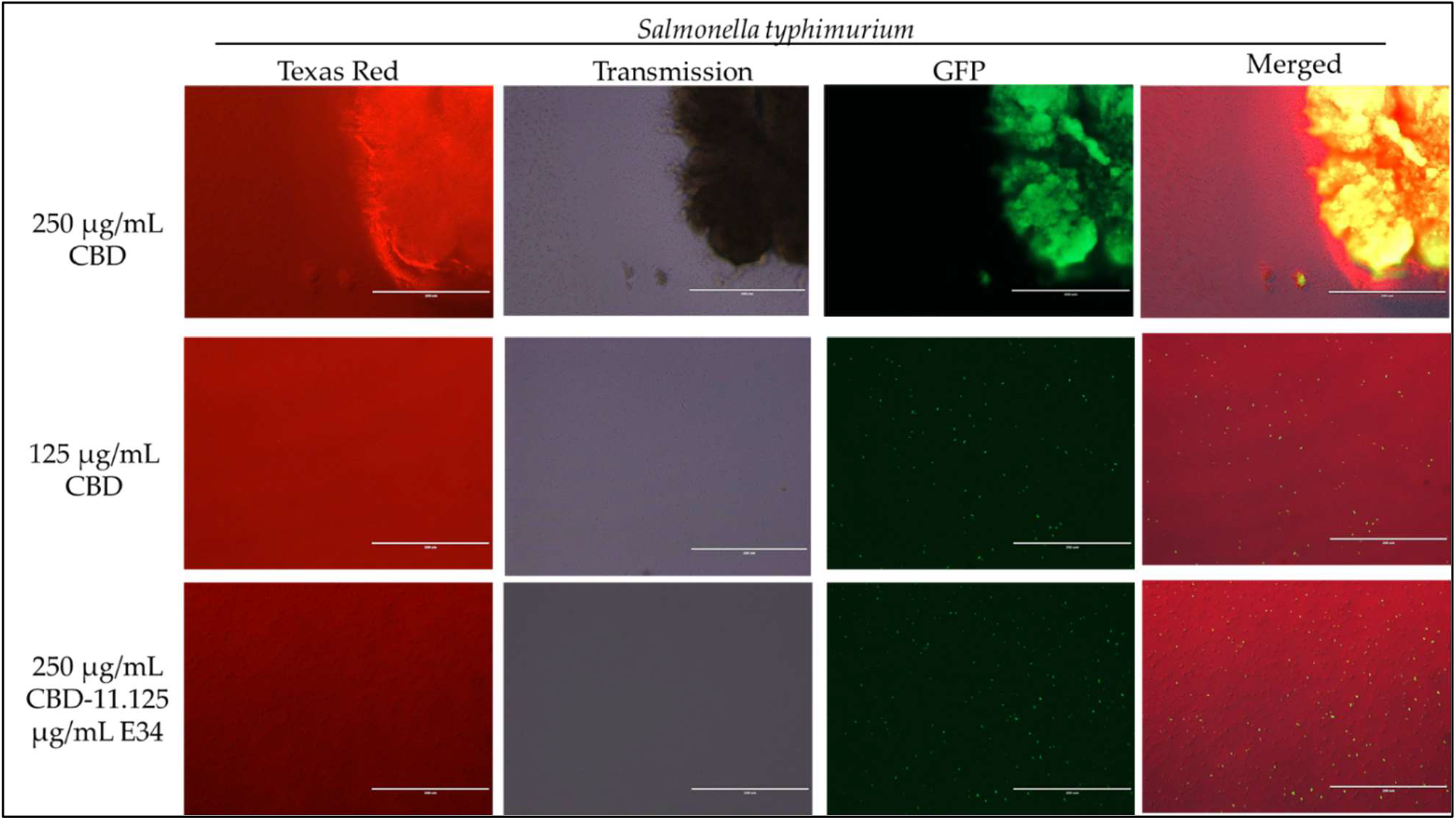
Immunofluorescence analysis of *S. typhimurium* (ST) at high concentrations of CBD (250 μg/mL, 125 μg/mL, and 250 μg/mL + 11.125 μg/mL E34 TSP) treatment. As shown in all panels, treatment at higher concentrations killed the *S typhimurium*. However, bacteria seemed to cluster together into biofilms enhancing their ability to survive (See top panel).

**Figure 7.**
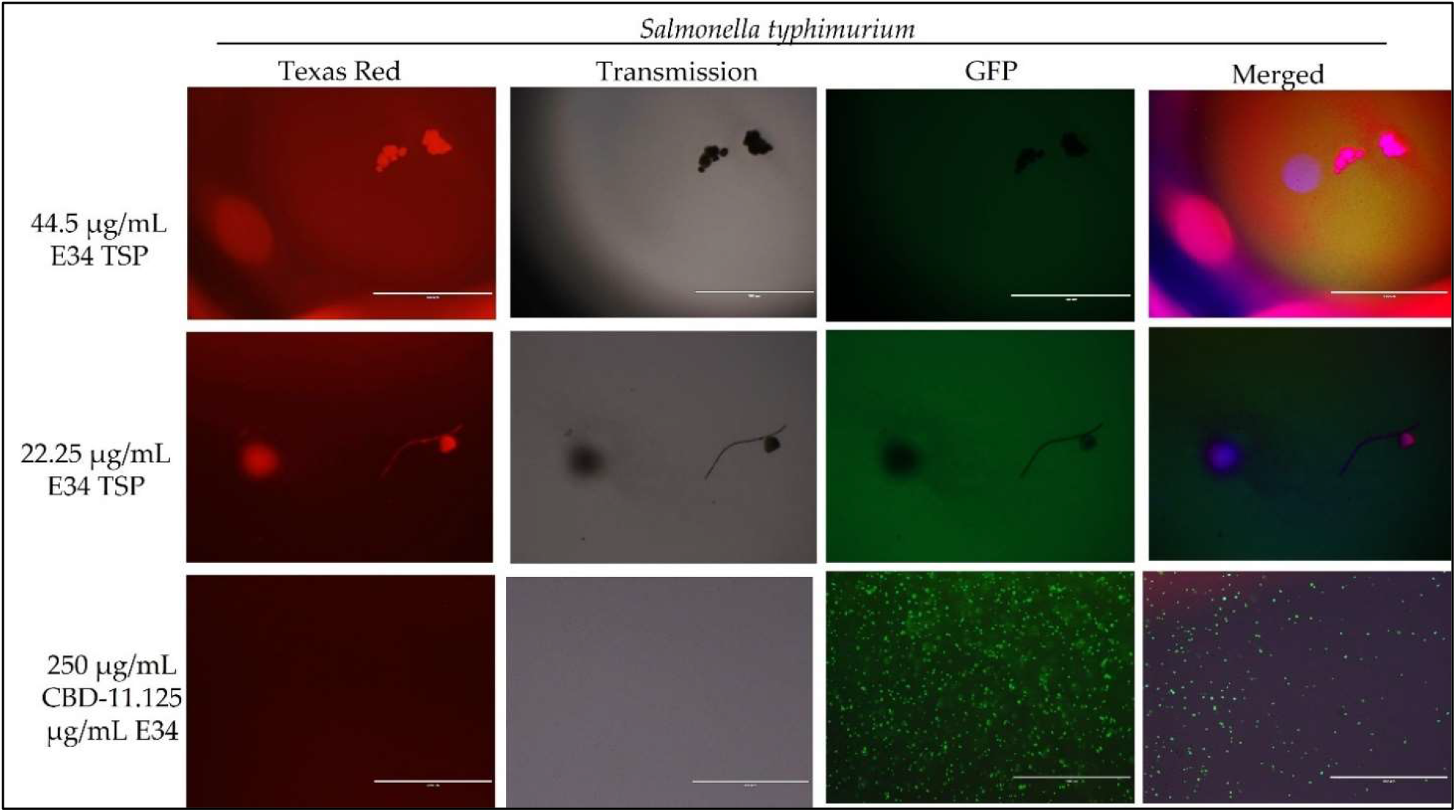
Immunofluorescence analysis of *S. typhimurium* (ST) at higher concentrations of E34 TSP; 44.5 μg/mL E34 TSP, 22.25 μg/mL E34 TSP, and the combination treatment of 250 μg/mL CBD + 11.125 μg/mL E34 TSP. As shown in all panels, treatment at higher concentrations killed the *S. typhimurium*. However, bacteria seemed to survive the combination treatment of CBD and E34 TSP especially at the higher concentration of 250 μg/mL CBD.

**Figure 8.**
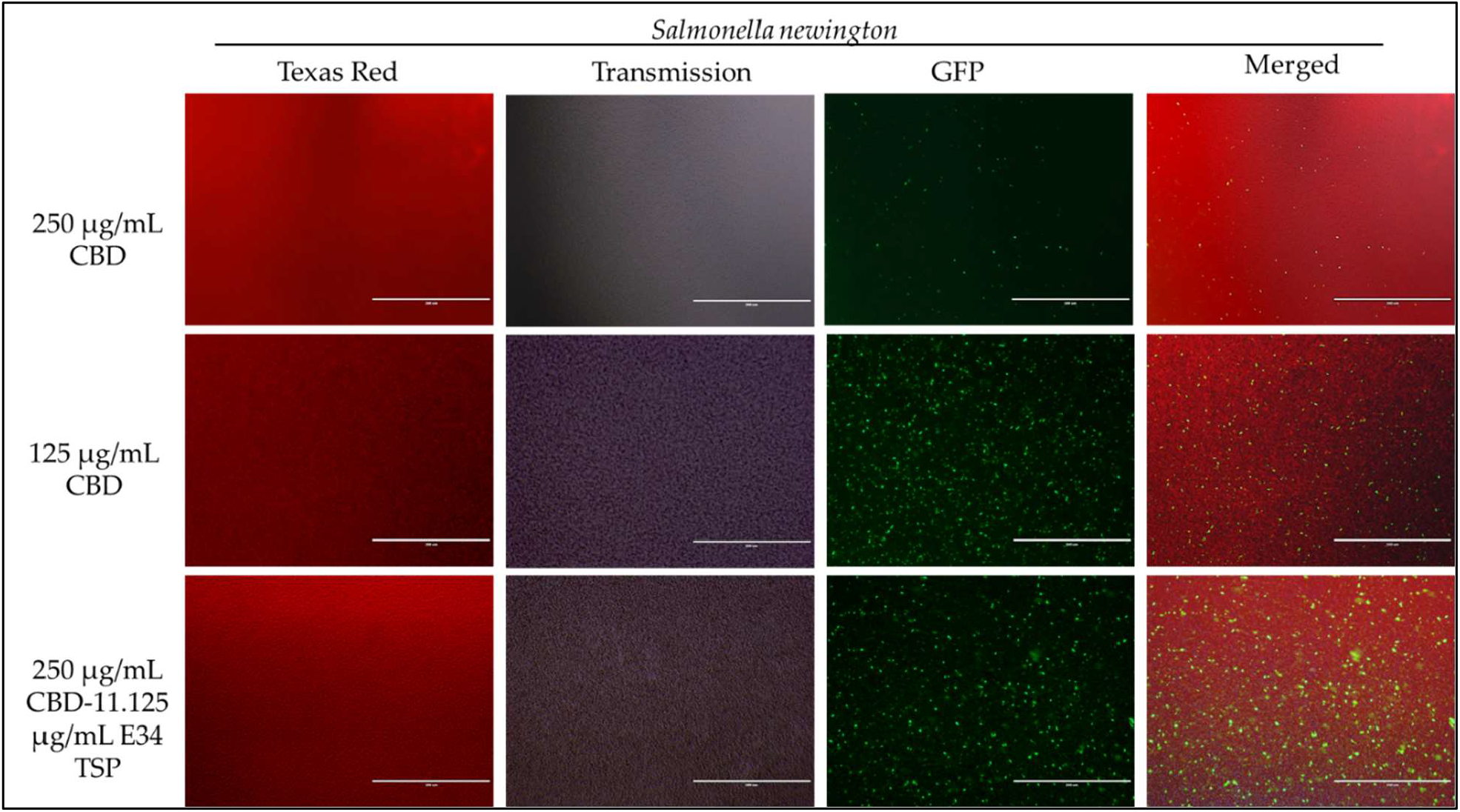
Immunofluorescence analysis of *S. newington* (SN) at higher concentrations of CBD (250 μg/mL, 125 μg/mL, and 250 μg/mL + 11.125 μg/mL E34 TSP) treatments. Generally, there is observed higher killing of *S. newington* in all treatments. Comparatively, there seems to be an observed lower killing of *S. newington* at the combination treatment.

**Figure 9.**
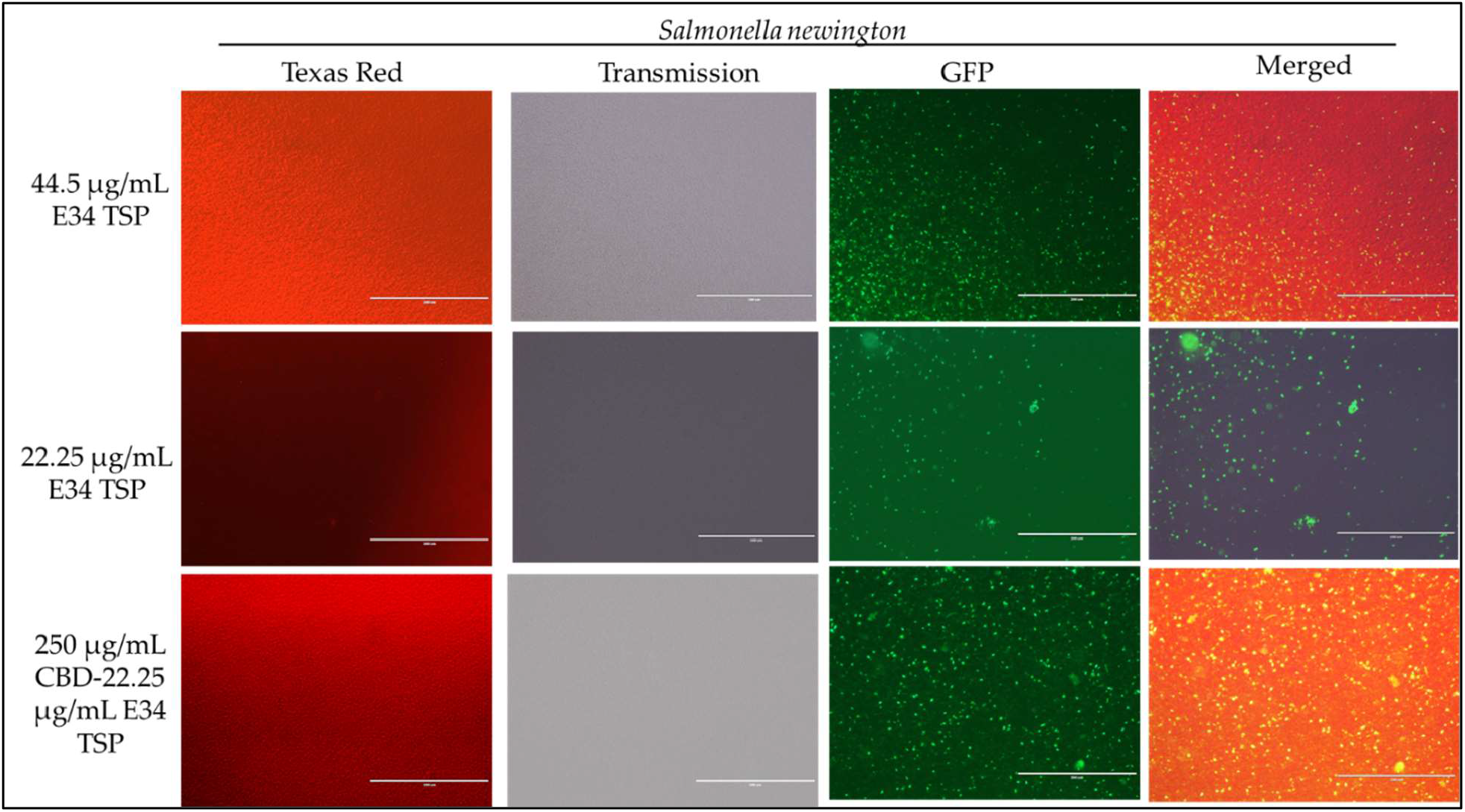
Immunofluorescence analysis of *S. newington* (SN) treated to low concentrations of E34 TSP; 44.5 μg/mL, 22.25 μg/mL, and a combination treatment of 250 μg/mL CBD + 22.25 μg/mL E34 TSP. The immunofluorescence images showed that the combination treatment (CBD and E34 TSP) had poorer membrane disruptive activities against *S. newington* as compared to the monotreatment with E34 TSP.

### 3.4 Time dependent analysis of E34 TSP and CBD treatments on *S. typhimurium* and *S. newington* growth at lag phase

To understand the effect of time kinetics of E34 TSP, CBD, or their combination treatment to *S. typhimurium* and *S. newington*, we subjected bacterial cells growing at lag phase (approximately 0.081 OD_600_) to E34 TSP and CBD treatment.

At the lag phase treatment, while all performed better than the control, treatment of *S. typhimurium* to 44.5 μg/mL and 22.25 μg/mL of E34 TSP performed significantly higher in inhibiting the bacterial growth. The combination treatment of 250 μg/mL CBD and 44.5 μg/mL E34 TSP also performed significantly better in inhibiting the *S. typhimurium* growth than the 250 μg/mL and 22.25 μg/mL combination treatment or the 250 μg/mL CBD monotreatment (**Figure 10A**). In **Figure 10B** however, only E34 TSP treatment at concentrations 22.25 μg/mL and 44.5 μg/mL showed high inhibition of *S. newington*, all other treatments showed poor inhibition kinetics. Comparatively, as shown in **Figure 10C**, the control group as expected showed the highest increase in OD_600_ of 38% compared to the other treatments, followed by the 250 μg/mL CBD and 22.25 μg/mL E34 TSP combination treatment which recorded 22%. Thus, indicating that these two treatments performed the poorest in inhibiting *S. typhimurium* growth at lag phase at the 12 h time point. The 22.25 μg/mL E34 TSP treatment showed the best inhibitory characteristic at the 12 h time point, recording 10%.

**Figure 10.**
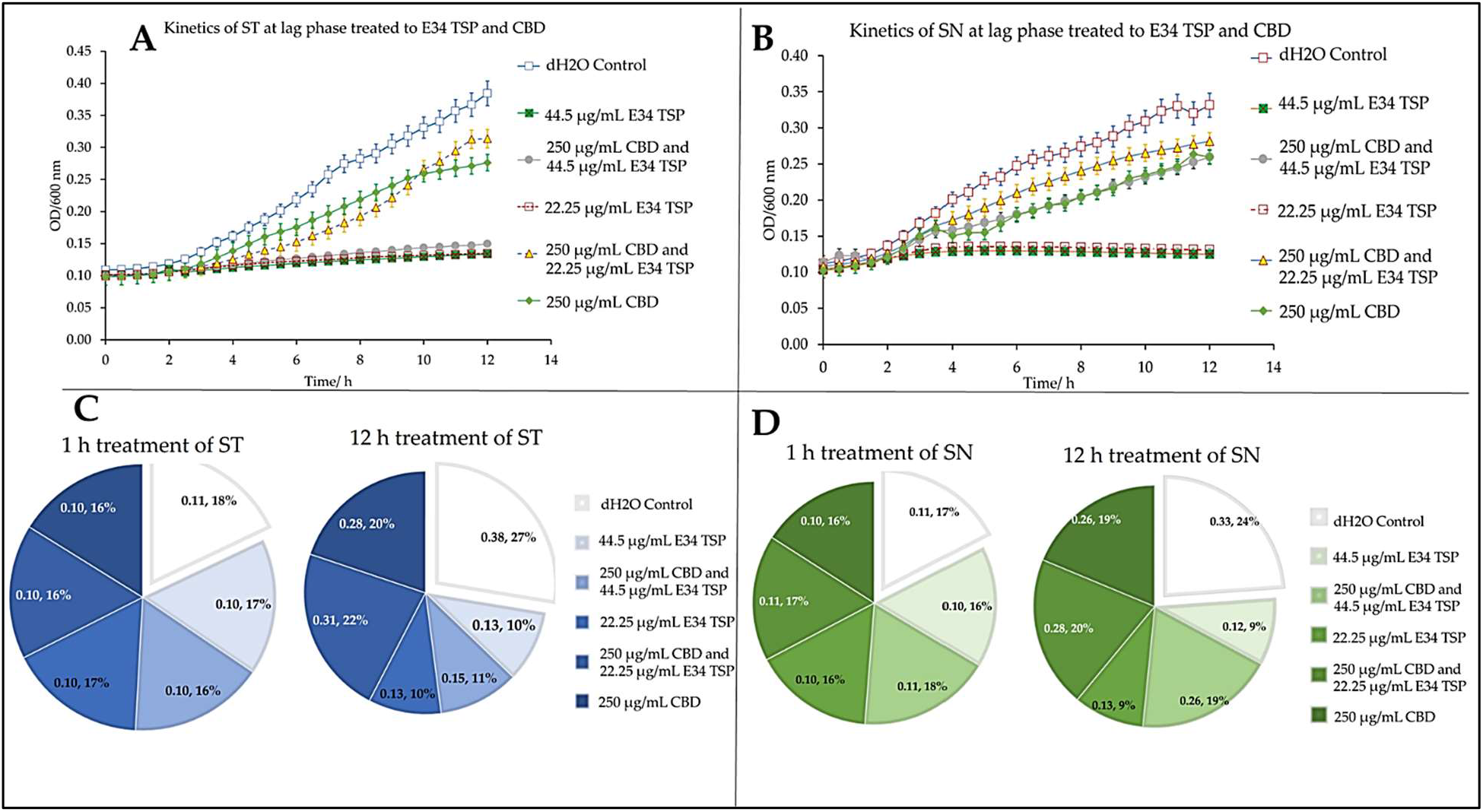
Kinetics of combined treatment of E34 TSP with CBD on **(A)** *S. typhimurium* (ST) in the lag growth phase, **(B)** *S. newington* (SN) in the lag growth phase. **(C)** Pie chart illustrating the relative percent changes in OD_600_ of *S. typhimurium* among treatments. **(D)** Pie chart illustrating the relative percent changes in OD_600_ of *S. newington* among treatments. Data shown are from three independent experiments expressed as means ± standard deviations.

In treating *S. newington*, as shown in **Figure 10D**, the best inhibitory effect again was observed in the 22.25 μg/mL E34 TSP treatment, which recorded a comparative rate of 16% at the 1 h time point and reduced this to 9% at the 12 h time point.

### 3.5 Dose dependent analysis of E34 TSP and CBD treatment on *S. typhimurium* and *S. newington*

To investigate how the concentration of E34 TSP, CBD or CBD-E34 TSP combinations could inhibit bacterial cell growth, we treated *S. typhimurium* and *S. newington* to varying concentrations of E34 TSP, CBD, and combinations of CBD-E34 TSP. **Figures 11 A-D** show the inhibition curves at the various treatment doses.

**Figure 11A.**
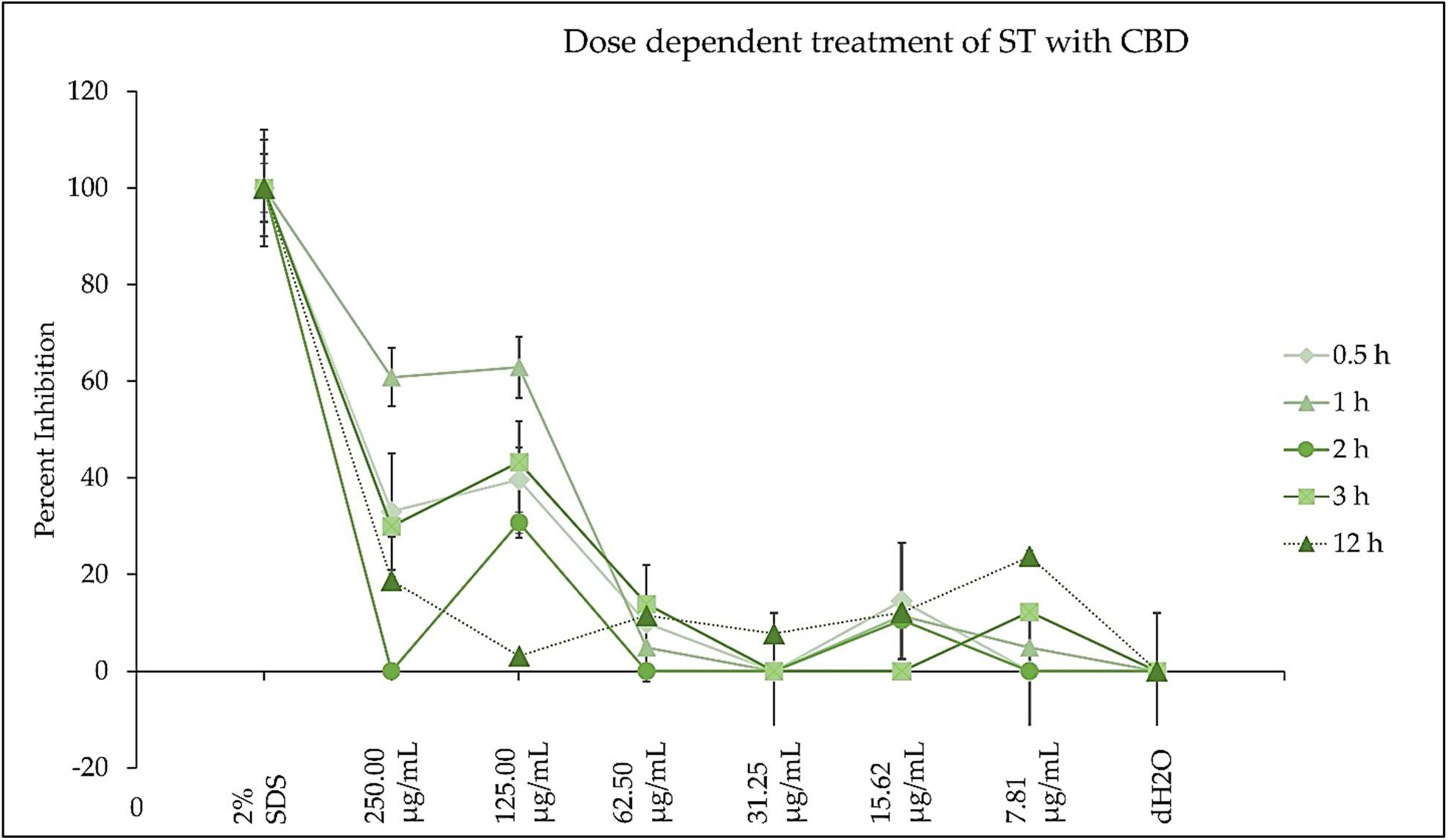
The effect of CBD on *S. typhimurium* presented as means ± standard deviation. Data shown are from three independent experiments expressed as means ± standard deviations. As can be seen, CBD although showed slightly higher inhibition at higher doses, it seemed not to exhibit time dependent inhibition of *S. newington*.

**Figure 11B.**
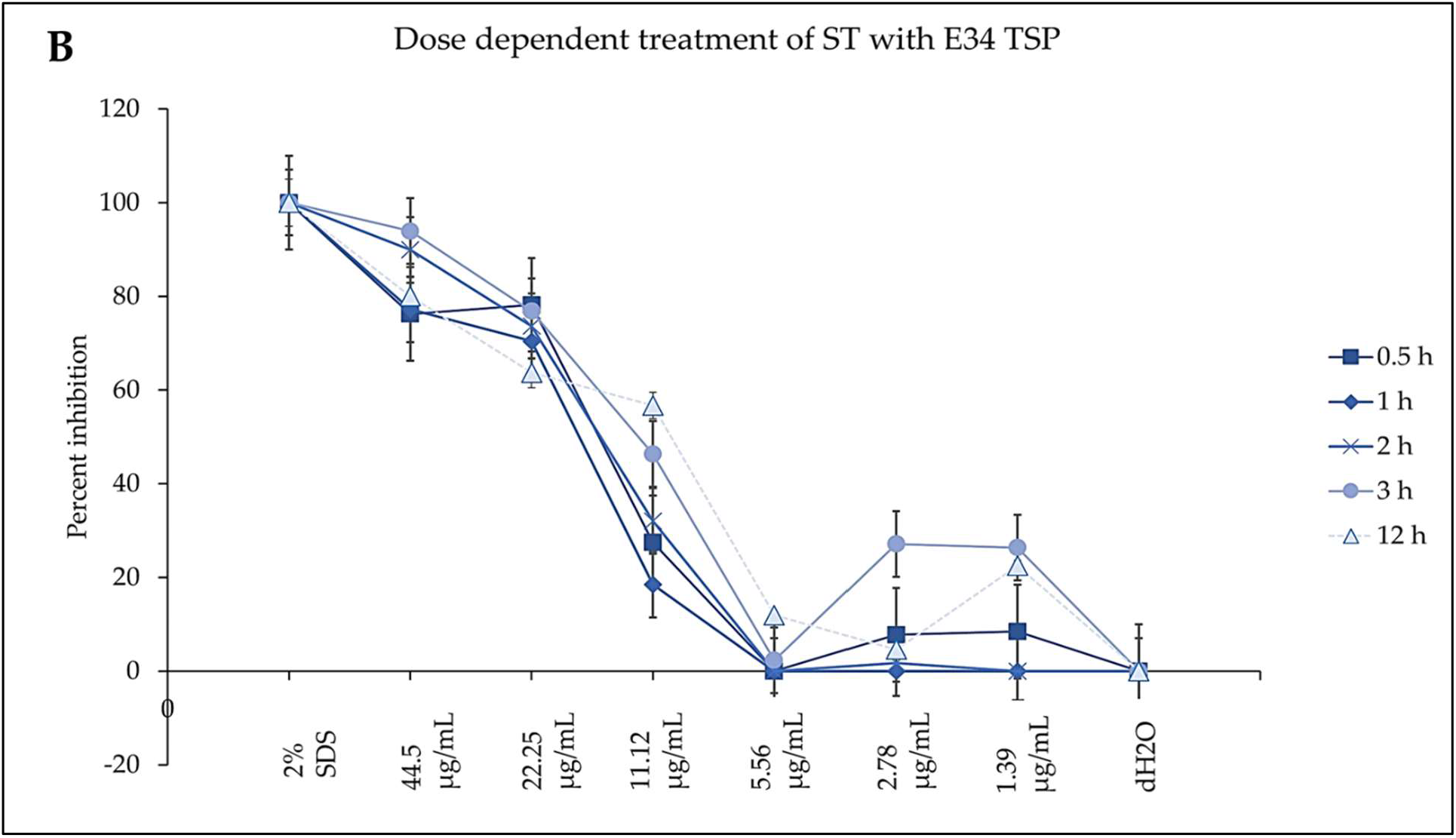
The effect of E34 TSP on *S. typhimurium* presented as means ± standard deviation. Data shown are from three independent experiments expressed as means ± standard deviations. As can be seen, E34 TSP seems to inhibit *S. typhimurium* in both time and dose dependent nature.

**Figure 11C.**
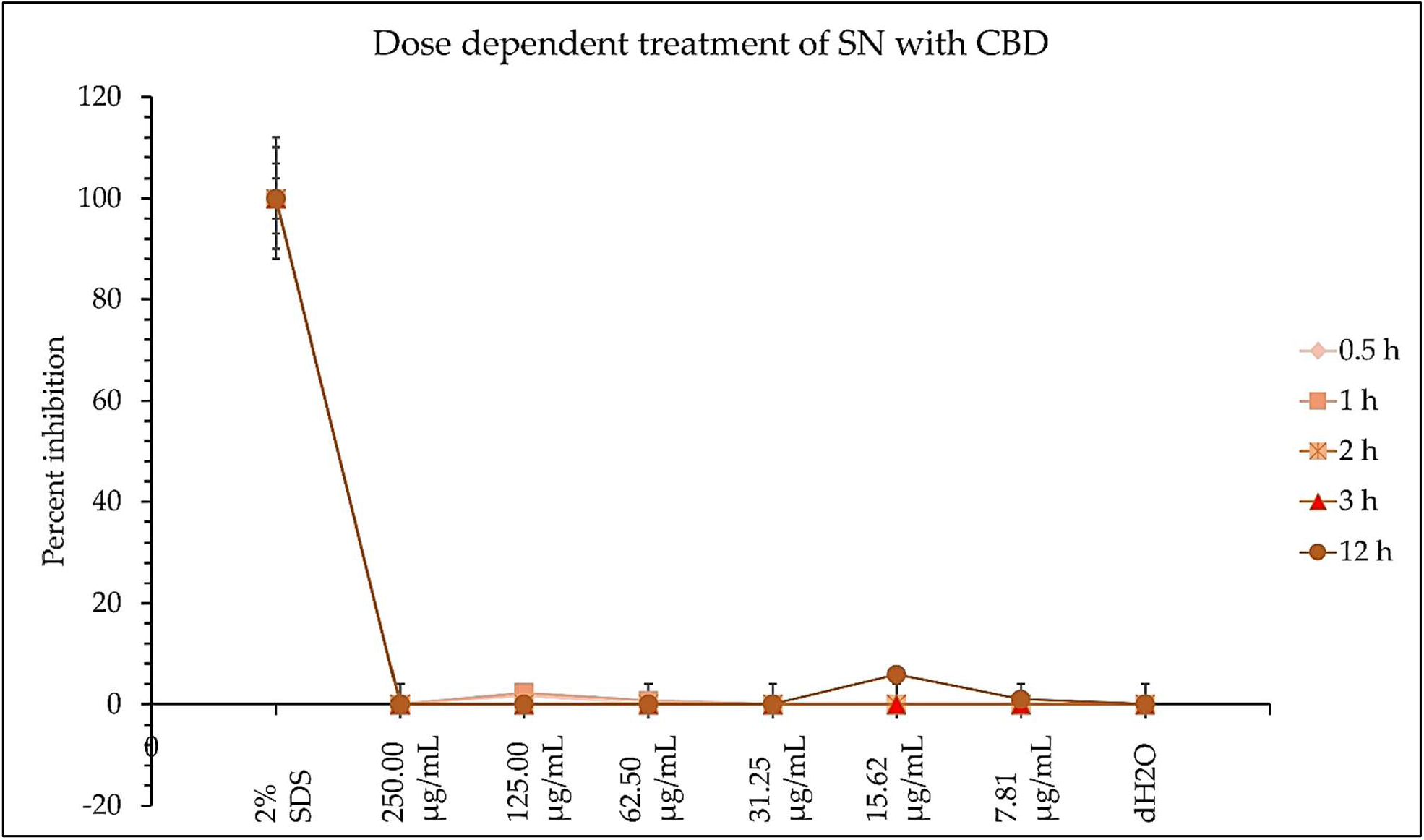
The effect of CBD on *S. newington* presented as means ± standard deviation. Data shown are from three independent experiments expressed as means ± standard deviations. As shown, CBD did not show any significant inhibition of *S. newington* in all treatments.

**Figure 11D.**
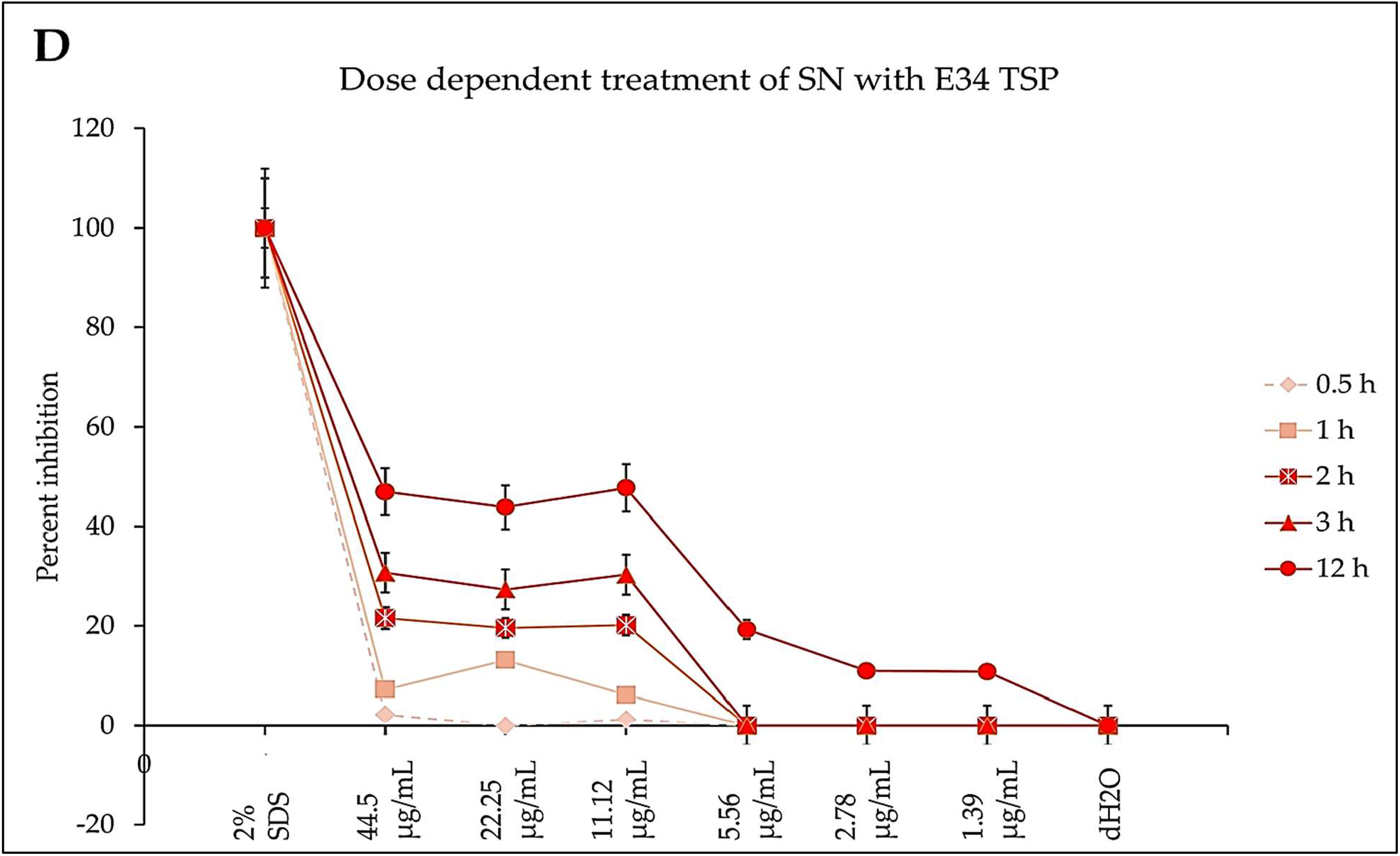
The effect of E34 TSP on *S. newington* presented as means ± standard deviation. Data shown are from three independent experiments expressed as means ± standard deviation. As can be observed, E34 TSP seems to inhibit *S. newington* in both time and dose dependent nature.

### 3.6 E34 TSP treatment potentiate bacterial membrane disruption

Given that there was observed substantial inhibition of *S. typhimurium* and *S. newington* caused by E34 TSP on the CBD-resistant strains, we found it important to investigate the possible mechanism of the bacterial killing mechanisms. We reasoned that it might be due to membrane integrity disruption. For this reason, we analyzed E34 TSP and CBD treated samples of CBD-resistant *S. typhimurium* using 0.8% agarose gel. Previous works by **Wassmann, et al., 2020** revealed that CBD is an effective helper compound in combination with bacitracin to kill Gram-positive bacteria via possible membrane disruption. Furthermore, **Gildea et al., 2022** also demonstrated the CBD in combination with polymyxin B was potent bactericidal agent whose mechanism of action was membrane lysis. In this study, treatment of CBD resistant strains of *S. typhimurium* to CBD produced relatively denser bands than the control group as indicated in Figure 12. The densities of the bands designated as non-migrated bacterial DNA showed highest in the Control group, followed by the CBD treatment group, E34 TSP treatment group showed the lowest density of bands in the non-migrated bacterial DNA. This points to the possibility that E34 TSP caused the highest bacterial membrane lysis, thus genomic content was free to migrate out of the cells. As depicted in the Migrated Bacterial DNA, the E34 TSP treated samples also gave the highest densities compared to both the control group and the CBD treated samples. Another interesting observation was the presence of short DNA fragments. These fragments were due to possible endonuclease activity during the treatment administration. As can be observed in **Figure 12**, most of these fragments are centered on the E34 TSP treated lanes, indicating further the possibility of E34 TSP exerting a much higher lytic activity on *S. typhimurium* than CBD. The inability of CBD to exert high lytic action on *S. typhimurium* could be due in part to the resistance of these strains to CBD. We infer that, the resistance mechanism is possibly via membrane content modifications that enforced lesser susceptibility to CBD interactions.

**Figure 12.**
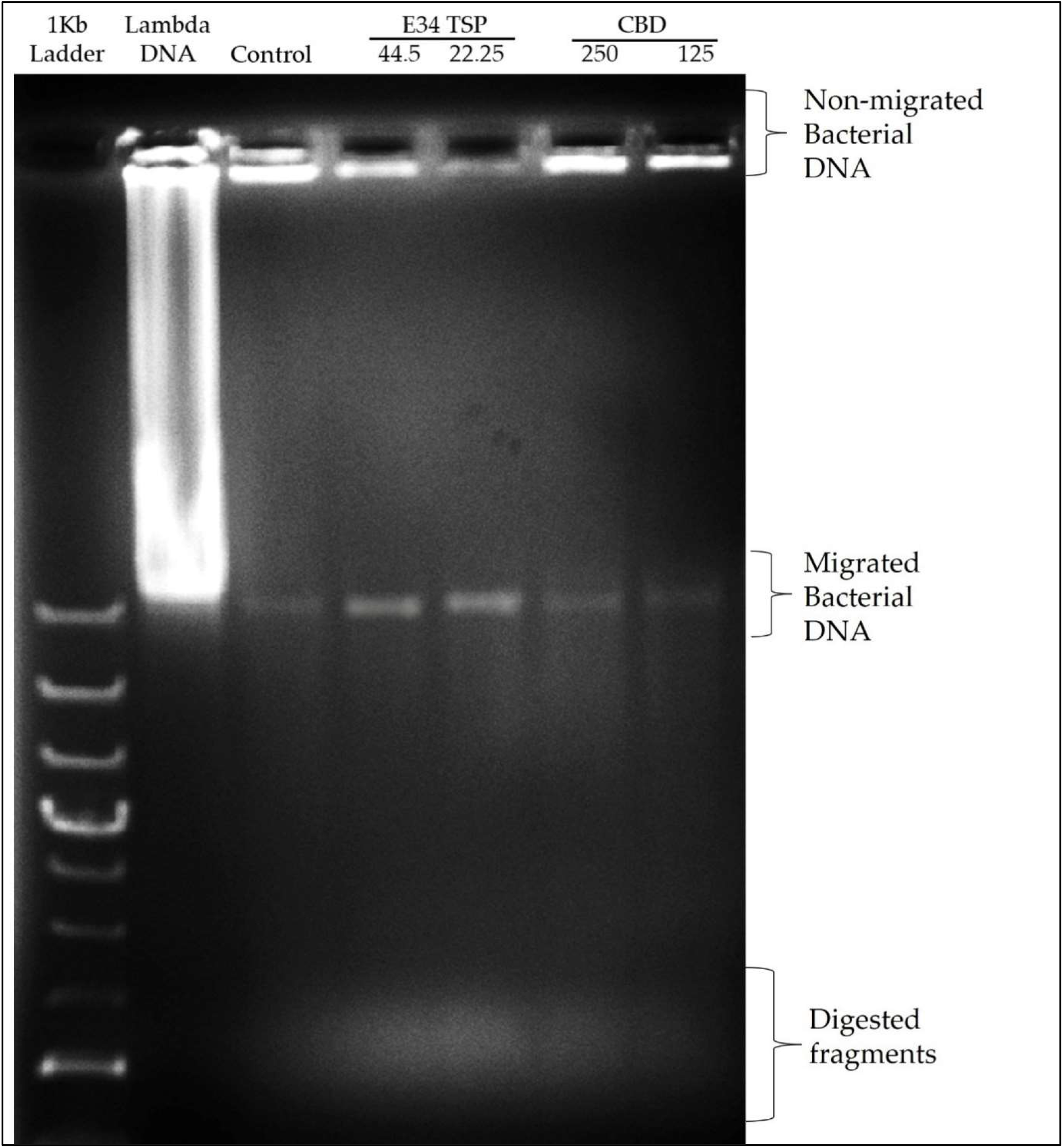
Agarose gel analysis of *S. typhimurium* treated to CBD, or E34 TSP. 30 μL of *S. typhimurium* samples treated to CBD or E34 TSP were loaded into each well, run at 90 volts for 1 h.

### 3.7 Analysis of dehydrogenase activity of treated *S. typhimurium* and *S. newington*

The glutamate dehydrogenase of *Salmonella* is one critical reductase that is modulated by various metabolites, which include amongst others the glycolytic TCA cycle and related intermediates, such as PEP, pyruvate, acetyl CoA, citrate, etc. Another set of metabolites that actively regulate *Salmonella* glutamate dehydrogenase activity are the purine nucleotides (AMP, ADP, ATP, CAMP, GMP, GDP, GTP). For instance, ATP is a potent activator of glutamate dehydrogenase (**Coulton and Kapoor, 1973**). Thus, to assess the effect of CBD treatment to glutamate dehydrogenase of *Salmonella* spp. used in this study, we employed resazurin assay.

For *S. typhimurium*, the effect of CBD on the bacteria dehydrogenase activity seems to negatively correlate with the concentration of CBD at the 1 h time point. E34 TSP showed no significant differences in all doses. The *S. newington* strain showed lower relative values when treated to E34 TSP at all concentrations except at 2.78 μg/mL and 1.39 μg/mL at the 1 h time point. CBD at all concentrations showed no significant effect on *S. newington* at 1 h time point. In general, 1 h posttreatment of *S. typhimurium* with CBD showed far lower dehydrogenase activity than E34 TSP treatments of the same strain. This is unexpected since our previous results demonstrated that E34 TSP showed higher killing of *S. typhimurium* than CBD (**Figure 4**). This might be due in part to the inability of E34 TSP to inhibit the dehydrogenase enzyme in *S. typhimurium* whereas CBD does inhibit it. A question remaining however, is why then we have less killing effect of CBD than E34 TSP in *S. typhimurium*. To answer this, we infer that, there might be a possible alternative dehydrogenase pathway utilized by *S. typhimurium* in energy metabolism. In treating *S. newington* however, E34 TSP treatments showed relatively lower dehydrogenase activity compared to the CBD treatment.

For *S. typhimurium*, the effect of CBD and E34 TSP was not hugely different from each other except the 31.25 μg/mL CBD treatment which considerably limited the dehydrogenase activity. The treatment of *S. newington* to E34 TSP however, showed general shutdown of the dehydrogenase enzyme at the 5 h time point. Interestingly, CBD treatment to the same strain produced slightly higher dehydrogenase activity as compared to the E34 TSP treatment. This trend is in consonance with the killing ability of E34 TSP on *S. newington* than CBD does. All SDS controls showed the lowest activity indicative of dead bacteria. Comparing the 5 h treatment to 1 h treatment, all the samples showed significantly lower dehydrogenase activity at 5 h than 1 h time point. This is indicative of the killing ability of all the treatments.

## 4.0 Discussion

Due to the emergence of multidrug-resistant bacteria, the use of bacteriophage as an alternative to antibiotics has been reconsidered. However, this approach can be very effective when combined with other agents. The combination of these two agents can provide better bacterial suppression and lower the chances of the development of resistance. It also allows for more efficient penetration into the bacterial population. Although neutral effects have been observed in some studies, combined approaches are still considered important in preventing the development of multidrug-resistant bacteria (**Barbu et al., 2016**). The main culprit for multi-drug resistance has been excessive use of antibiotics, which jeopardizes their effectiveness for controlling these pathogenic agents (**Tagliaferri et al., 2019**). Considering this, bacteriophages and their derivatives are becoming more widely accepted as viable complementary techniques for use in food safety, health, and medicine.

In this study, to investigate the possibility of using a phage protein (E34 TSP) in conjunction with CBD to treat CBD resistant strains of *Salmonella*, we expressed E34 TSP from BL21/DE3 cells using pET30a-LIC-E34 TSP, a clone which we previously created (**Ayariga et al., 2021**). E34 phage is a bacteriophage that infects *S. newington*, and it uses its hydrolase machinery which is the tailspike protein (TSP) for the initial interaction and anchoring of the phage particle to the LPS of the cell and subsequent hydrolysis and anchoring of the LPS to the membrane of the bacteria. In this study, the E34 TSP was expressed under the control of the T7 promoter in pET30a-LIC vector, purified and combined with CBD as an antibacterial agent against CBD-resistant strains of *Salmonella*. Validation of the cloned insert was achieved via PCR reaction using two primers that amplified exactly the tailspike gene, gp19 in the clone. An agarose gel electrophoresis of PCR product demonstrated that it carried the exact size of 1.818 kbp insert (**Figure 1A**). Non-recombinants control did not yield any band, indicative of the absence of the insert. Cobalt-NTA column was used for affinity purification via FPLC (chromatograph shown in **Figure 2**), and samples were run on SDS PAGE to validate the purity of the protein.

The ability of CBD to potently inhibit *S. typhimurium* was previously demonstrated in our laboratory (**Gildea et al., 2022**), other works by **Wassmann et al., 2020** also revealed that CBD is a novel antibiotic adjuvant that potentiates the effect of the bacitracin against Gram-positive bacteria (e.g., *Staphylococcus aureus, Listeria monocytogenes, and Enterococcus faecalis*). The same group in another study explored the mechanism of resistance to CBD by *Staphylococcus aureus*. They discovered through resistant strains genome sequencing that the farE/farR system encoding a fatty acid efflux pump (FarE) and its regulator (FarR) were mutated, which showed diminished susceptibility to both CBD and bacitracin (**Wassmann et al., 2022**).

Proceeding from lag phase, bacteria enter log phase, a phase characterized by exponential growth. One major biological characteristic of this phase is the high metabolic activities occurring, which is due in part to high DNA replication, RNA translation, cell wall biosynthesis, and in part due to high cell division. While bacteria are metabolically hyperactive at this phase, they are also most vulnerable too, it is in this phase that antibiotics and other antibacterial agents can produce their highest potency. Usually, most of these agents target bacteria cell wall synthesis (e.g. Beta-lactams), or protein synthesis (e.g. thermorubin, (**Bulkley et al., 2012**)), DNA transcription (e.g. Anthracycline**s** (**Bilardi et al, 2012**)), and RNA translation (e.g. spectinomycin (**Borovinskaya et al., 2007**)). In this work, we investigated the effect of treating *S. newington* and *S. typhimurium* to CBD, E34 TSP, and the combination of the two agents at both early and late log phase of the bacteria. As shown in **Figure 4** while generally all treatments performed better in *S. newington* than in *S. typhimurium*, the E34 TSP treatments gave the best inhibition than both CBD and CBD-E34 TSP combination. While it is predictable that E34 TSP monotreatment will perform better than CBD in reducing the bacterial growth, since these strains were CBD resistant, it is surprising that the combination treatment performed poorly. While these results necessitate a multiple dose analysis of CBD, and E34 TSP combinations, it seems safe to infer that there is probably an antagonistic interaction between the CBD and E34 TSP. This might be due in part to CBD probably binding to the catalytic site of the E34 TSP, thus blocking it from endorhamnosidase activity. Since these strains are CBD-resistant, inactivated E34 TSP and abundant CBD will have minimal effect on the bacteria. **Figures 5-9** depict the immunofluorescent images showing the ability of E34 TSP to kill *S. typhimurium* and *S. newington*. However, as shown in **Figure 6**, the bacteria seemed to pool into micro clusters in high CBD concentrations that possibly provided unique shields against the CBD. This clustering might enable the development of biofilm by the bacteria, thus enhancing their CBD resistance (**Horstmann et al., 2021**). Micro clustering enables quicker cell-cell communication in the form of quorum sensing or membrane vesicle trafficking. The membrane vesicles in bacteria play a role in cell-cell communication between bacteria themselves and between bacteria their hosts. Membrane vesicles are an important component of anti-bacterial resistance and thus have gained relevance in antibacterial resistance research. Given this, **Kosgodage et al., 2019** revealed that CBD affected the membrane vesicle profile and membrane vesicle release of bacteria. They reported that CBD had a strong inhibitory influence on membrane vesicle release of Gram-negative bacteria such as *E. coli* VCS257, and negligible inhibitory influence of CBD on Gram-positive bacteria such as *S. aureus* subsp. *aureus* Rosenbach 1884 membrane vesicle release. Thus, the micro clustering might be a defensive maneuver to overcome CBD anti-vesicle release property.

The antibacterial properties of CBD cannot be jettisoned. This is in the light of the fact that several studies have idealized the concept of utilizing this compound as an antimicrobial. The commencement of disease symptoms mostly starts after the lag phase of bacterial infection. This initial phase of the bacterial growth is characterized by cellular activity that typically involves the synthesis of proteins without much growth. To understand the effect of CBD on our CBD-resistant strains growing at lag phase, we treated *S. typhimurium* and *S. newington* to CBD, E34 TSP and CBD-E34 TSP combination. As it is well known, CBD is recognized as a potent antimicrobial. **Blaskovich et al., 2021** studied the antimicrobial characteristics of CBD-resistance in *Staphylococcus aureus, Streptococcus pneumonia*, and *Clostridioides difficile*. Their findings suggested that CBD has an impeccable impact on biofilm and topical *in vivo* efficacy. In this study, at the lag phase, while all treatments performed better than the control, monotreatment of *S. typhimurium* and *S. newington* to 44.5 μg/mL and 22.25 μg/mL of E34 TSP performed significantly higher in inhibiting the bacterial growth. The results corroborated with the data obtained in the log phase treatment indicating that bacteria growth phase did not significantly affect the performances of the treatments.

To understand the characteristic effect of our bacteria under different concentrations of E34 TSP and CBD at mid-log phase of the bacteria, we carried out dose dependent treatment of E34 TSP and CBD treatment on *S. typhimurium* and *S. newington*. As shown in **Figure 11A**, CBD showed slightly higher inhibition at higher doses, but did not exhibit time dependent inhibition of *S. newington*. E34 TSP demonstrated both time and dose dependent inhibition of *S. typhimurium* (**Figure 11B**). As expected, CBD failed to show any significant inhibition of *S. newington* in all treatments (**Figure 11C**). Treatment of *S. newington* to E34 TSP showed time and dose dependent inhibition (**Figure 11D**).

Also highlighted in this study is the membrane disruption potential of E34 TSP, CBD, and their combinations. CBD has been demonstrated to show potent membrane disruption as its primary mechanism of attack (**Blaskovich et al., 2021**). Intuitively, the same study also hinted for the first time how CBD deactivates the “urgent threat” pathogen *Neisseria gonorrhoeae*. Similarly, **Abichabki et al., 2022**, reported how CBD extract was effective against *Neisseria gonorrhoeae, Neisseria meningitides, Moraxella catarrhalis*, and *Mycobacterium tuberculosis*. Their CBD formulation, however, was in combination with polymyxin B which is a known antibiotic that shows very potent membrane disruption activity (**Grau-Campistany et al., 2016**). They stated that the activities were achieved at a polymyxin B concentration of ≤ 2 μg/mL and CBD concentration of ≤ 4 μg/mL, especially against *Klebsiella pneumonia, Escherichia coli*, and *Acinetobacter baumannii*. In our research work, we showed that CBD caused very slight membrane disruption compared to E34 TSP which caused a much higher membrane disruption (**Figure 12**). The lower membrane disruption of CBD might be attributable to the resistant strains used for the study.

In bacteria, respiratory chains are composed of various types of transport constituents, such as cytochromes, quinones, iron-sulfur proteins, and flavoproteins. The differential transport of protons and electrons through the cytoplasm leads to the formation of a proton gradient membrane. This membrane can be used to drive the formation of ATP through the F1/F0 ATPase (**Ingledew and Poole, 1984, Syroeshkin et al., 1998, Russell**, **2007**). Electrons are transported through a variety of redox carriers to reach the oxygen-rich environment during respiration. They can also be obtained from alternative terminal electron acceptors when oxygen is unavailable (**Tielens and Hellemond, 1998**). The glutamate dehydrogenase of *Salmonella* is one critical reductase that is modulated by various metabolites, which include amongst others the glycolytic, TCA cycle and related intermediates, such as PEP, pyruvate, acetyl CoA, citrate, etc. Another set of metabolites that actively regulate *Salmonella* glutamate dehydrogenase activity are the purine nucleotides (AMP, ADP, ATP, CAMP, GMP, GDP, GTP). For instance, ATP is a potent activator of glutamate dehydrogenase (**Coulton and Kapoor, 1973**). Thus, to assess the effect of CBD treatment to glutamate dehydrogenase of *Salmonella* spp. used in this study, we employed resazurin assay. As shown in **Figure 13A**, the effect of CBD on *S. typhimurium* dehydrogenase activity seems to negatively correlate with the concentration of CBD at the 1 h time point whereas E34 TSP showed no significant difference in all doses. In general, 1 h posttreatment of *S. typhimurium* with CBD showed far lower dehydrogenase activity than E34 TSP treatments of the same strain. This is unexpected since our previous results demonstrated that E34 TSP showed higher killing of *S. typhimurium* than CBD (**Figure 4**). This might be explained in part by proposing that E34 TSP does not interact with dehydrogenase enzyme in *S. typhimurium* whereas CBD does possibly interact with the enzyme. The concern, however, is why the less killing effect of CBD observed in *S. typhimurium* than E34 TSP. To answer this, we infer that there might be a possible alternative dehydrogenase pathway utilized by *S. typhimurium* in energy metabolism especially at the 1 h time point. In treating *S. newington* however, E34 TSP treatments showed relatively lower dehydrogenase activity compared to the CBD treatment.

**Figure 13A.**
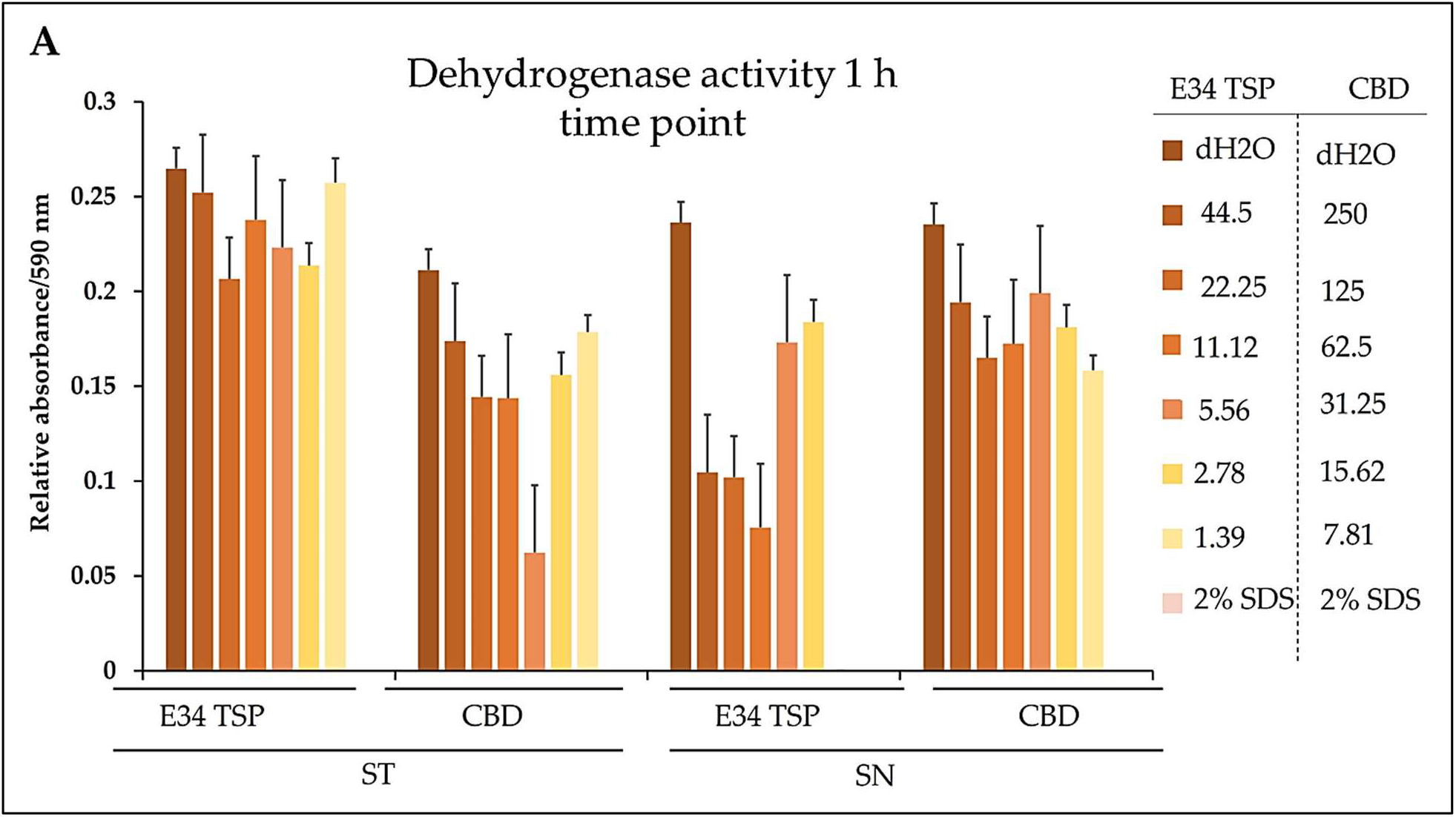
Chart illustrating the *S. typhimurium* and *S. newington* dehydrogenase activity in 1 h after treatment to varying concentrations of CBD or E34 TSP. Data shown are from three independent experiments expressed as means ± standard deviations.

**Figure 13B.**
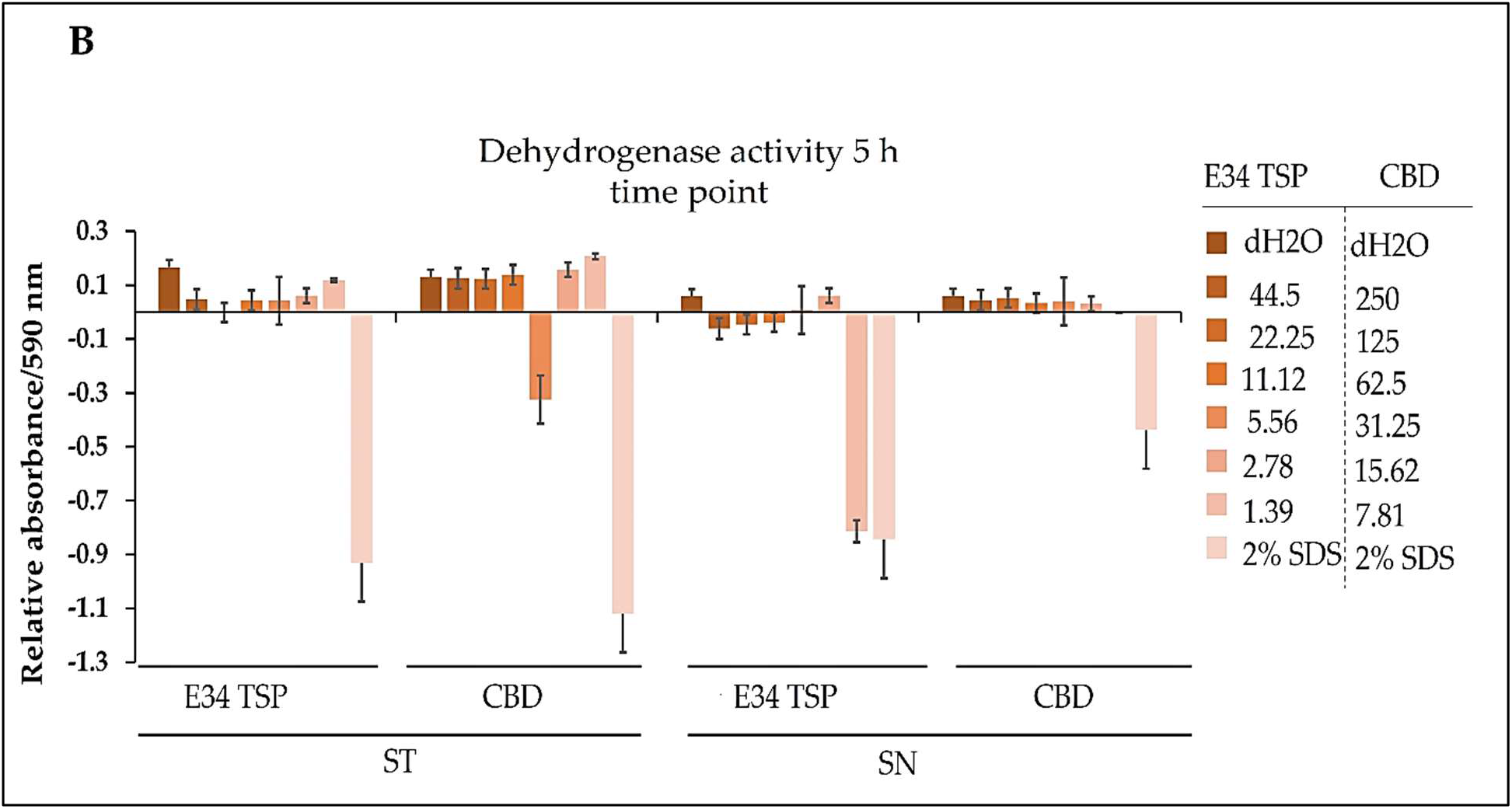
Chart illustrating the *S. typhimurium* and *S. newington* dehydrogenase activities in 5 h after treatment to varying concentrations of CBD, or E34 TSP. Data shown are from three independent experiments expressed as means ± standard deviations.

As shown in **Figure 13B**, treatment of *S. typhimurium* and *S. newington* showed a general decrease in dehydrogenase activity in 5 h. The most pronounced decrease however was observed in *S. newington* treated to E34 TSP which demonstrated a complete inactivation of the enzyme at the 5 h time point. Interestingly, CBD treatment to the same strain produced slightly higher dehydrogenase activity as compared to the E34 TSP treatment. This trend is in consonance with the killing ability of E34 TSP on *S. newington* than CBD does. All SDS controls showed the lowest activity indicative of dead bacteria. Comparing the 5 h treatment to 1 h treatment, all the samples showed significantly lower dehydrogenase activity at 5 h than 1 h time point. This is indicative of the killing ability of all the treatments.

## Conclusion

Given the rapid antimicrobial resistance development, it is crucial to look for other avenues to treat antimicrobial resistant bacteria. Our work demonstrated that even strains that are resistant to the potent CBD could still be inhibited by a phage protein such as E34 TSP. We also showed that the inhibition of the bacteria by E34 TSP was due in part to membrane disruption, and dehydrogenase inactivation by the protein. A CBD-E34 TSP combination treatment resulted in lower killing ability of the treatment. This interesting finding indicates that CBD might have an antagonistic property against the protein. We proposed that CBD binds to the catalytic site of the E34 TSP hydrolase thus hindering the protein’s endorhamnosidase activity on the bacteria. Further research is needed to fully elucidate the mechanism of inactivation of E34 TSP by CBD. Moreso, it will be very interesting to explore the genetic basis of the resistance development of these two strains of bacteria to CBD. In conclusion, this work highlights the crucial role phage protein such as E34 TSP could play in pathogenic bacterial control.

## Author Contributions

Conceptualization JAA; methodology, II and JAA; software, II and JAA; validation, JAA, OSA, and MS; formal analysis, JAA, JX, MS, and OSA; investigation, II, JAA, AA, JX, MS, and OSA; resources, JAA, MS, and OSA; data curation, JAA, JX, and OSA.; writing—original draft preparation, II, and JAA; writing—review and editing, II, JAA, JX, OSA; visualization, II and JAA; supervision, JAA, and OSA; project administration, OSA; funding acquisition, OSA. All authors have read and agreed to the published version of the manuscript.

## Informed Consent Statement

Not applicable.

## Data Availability Statement

Data is contained within the article.

## Acknowledgments

The authors acknowledge late Professor Robert Villafane for mentorship. We also acknowledge Alabama State University, C-STEM for supplies and Laboratory space. The authors acknowledge receiving funding from United States Department of Education, Title III-HBGI-RES.

## Sample Availability

Samples of the compounds are available from the authors.

## References

1. Abichabki, N., Zacharias, L.V., Moreira, N.C., Bellissimo-Rodrigues, F., Moreira, F.L., Benzi, J.R., Ogasawara, T., Ferreira, J.C., Ribeiro, C.M., Pavan, F.R. and Pereira, L.R., 2022. Potential cannabidiol (CBD) repurposing as antibacterial and promising therapy of CBD plus polymyxin B (PB) against PB-resistant gram-negative bacilli. Scientific reports, 12(1), pp.1–15.

2. Adams, R., Pease, D.C., Cain, C.K. and Clark, J.H., 1940. Structure of cannabidiol. VI. Isomerization of cannabidiol to tetrahydrocannabinol, a physiologically active product. Conversion of cannabidiol to cannabinol1. Journal of the American Chemical Society, 62(9), pp.2402–2405.

3. Alonso, B., Cruces, R., Pérez, A., Sánchez-Carrillo, C. and Guembe, M., 2017. Comparison of the XTT and resazurin assays for quantification of the metabolic activity of Staphylococcus aureus biofilm. Journal of Microbiological Methods, 139, pp.135–137.

4. Ayariga, J.A., Abugri, D.A., Amrutha, B. and Villafane, R., 2022. Capsaicin potently blocks Salmonella typhimurium invasion of Vero cells. Antibiotics, 11(5), p.666.

5. Ayariga, J.A.; Gildea, L.; Villafane, R. ε34 phage tailspike protein is resistant to trypsin and inhibits Salmonella biofilm formation. Enliven Microb. Microb. Tech. 2022, 9, 002.

6. Azeredo, J. and Sutherland, I.W., 2008. The use of phages for the removal of infectious biofilms. Current pharmaceutical biotechnology, 9(4), pp.261–266.

7. Balasubramanian, R., Im, J., Lee, J.S., Jeon, H.J., Mogeni, O.D., Kim, J.H., Rakotozandrindrainy, R., Baker, S. and Marks, F., 2019. The global burden and epidemiology of invasive non-typhoidal Salmonella infections. Human vaccines & immunotherapeutics, 15(6), pp.1421–1426.

8. Barbu, E.M., Cady, K.C. and Hubby, B., 2016. Phage therapy in the era of synthetic biology. Cold Spring Harbor perspectives in biology, 8(10), p. a023879.

9. Bartner, L.R., McGrath, S., Rao, S., Hyatt, L.K. and Wittenburg, L.A., 2018. Pharmacokinetics of cannabidiol administered by 3 delivery methods at 2 different dosages to healthy dogs. Canadian Journal of Veterinary Research, 82(3), pp.178–183.

10. Bilardi, R.A., Kimura, K.I., Phillips, D.R. and Cutts, S.M., 2012. Processing of anthracycline-DNA adducts via DNA replication and interstrand crosslink repair pathways. Biochemical pharmacology, 83(9), pp.1241–1250.

11. Blaskovich, M.A., Kavanagh, A.M., Elliott, A.G., Zhang, B., Ramu, S., Amado, M., Lowe, G.J., Hinton, A.O., Pham, D.M.T., Zuegg, J. and Beare, N., 2021. The antimicrobial potential of cannabidiol. Communications biology, 4(1), pp.1–18.

12. Borovinskaya, M.A., Shoji, S., Holton, J.M., Fredrick, K. and Cate, J.H., 2007. A steric block in translation caused by the antibiotic spectinomycin. ACS chemical biology, 2(8), pp.545–552.

13. Borysowski, J.; Weber-Dąbrowska, B.; Górski, A. Bacteriophage endolysins as a novel class of antibacterial agents. Exp. Biol. Med. 2006, 231, 366–377. [Google Scholar] [CrossRef].

14. Britch, S.C., Babalonis, S. and Walsh, S.L., 2021. Cannabidiol: pharmacology and therapeutic targets. Psychopharmacology, 238(1), pp.9–28.

15. Brunetti, P., Faro, A.F.L., Pirani, F., Berretta, P., Pacifici, R., Pichini, S. and Busardò, F.P., 2020. Pharmacology and legal status of cannabidiol. Annali dell’Istituto Superiore di Sanità, 56(3), pp.285–291.

16. Bulkley, D., Johnson, F. and Steitz, T.A., 2012. The antibiotic thermorubin inhibits protein synthesis by binding to inter-subunit bridge B2a of the ribosome. Journal of molecular biology, 416(4), pp.571–578.

17. Chan, J.Z. and Duncan, R.E., 2021. Regulatory effects of cannabidiol on mitochondrial functions: a review. Cells, 10(5), p.1251.

18. Chanishvili N. Phage therapy--history from Twort and d’Herelle through Soviet experience to current approaches. Adv Virus Res. 2012; 83:3–40. doi: 10.1016/B978-0-12-394438-2.00001-3. PMID: 22748807.

19. Clokie, M.R., Millard, A.D., Letarov, A.V. and Heaphy, S., 2011. Phages in nature. Bacteriophage, 1(1), pp.31–45.

20. Coulton, J.W. and Kapoor, M., 1973. Studies on the kinetics and regulation of glutamate dehydrogenase of Salmonella typhimurium. Canadian Journal of Microbiology, 19(4), pp.439–450.

21. Fernandes, M.F., Chan, J.Z., Hung, C.C.J., Tomczewski, M.V. and Duncan, R.E., 2022. Effect of cannabidiol on apoptosis and cellular interferon and interferon-stimulated gene responses to the SARS-CoV-2 genes ORF8, ORF10 and M protein. Life Sciences, p.120624.

22. Fernandes, M.F., Chan, J.Z., Hung, C.C.J., Tomczewski, M.V. and Duncan, R.E., 2022. Effect of cannabidiol on apoptosis and cellular interferon and interferon-stimulated gene responses to the SARS-CoV-2 genes ORF8, ORF10 and M protein. Life Sciences, p.120624.

23. Fischetti, V.A. Bacteriophage lytic enzymes: Novel anti-infectives. Trends Microbiol. 2005, 13, 491–496. [Google Scholar] [CrossRef].

24. Gildea, L., Ayariga, J.A., Ajayi, O.S., Xu, J., Villafane, R. and Samuel-Foo, M., 2022. Cannabis sativa CBD Extract Shows Promising Antibacterial Activity against Salmonella typhimurium and S. newington. Molecules, 27(9), p.2669.

25. Gildea, L.; Ayariga, J.; Xu, J.; Villafane, R.; Boakai, R.; Samuel-Foo, M.; Ajayi, O. Cannabis sativa CBD Extract Exhibits Synergy with Broad-Spectrum Antibiotics against Salmonella typhimurium. Preprints 2022, 2022090143 (doi: 10.20944/preprints202209.0143.v1).

26. Grau-Campistany, A., Manresa, Á., Pujol, M., Rabanal, F. and Cajal, Y., 2016. Tryptophan-containing lipopeptide antibiotics derived from polymyxin B with activity against Gram positive and Gram-negative bacteria. Biochimica et Biophysica Acta (BBA)-Biomembranes, 1858(2), pp.333–343.

27. Gut, A.M., Vasiljevic, T., Yeager, T. and Donkor, O.N., 2018. Salmonella infection–prevention and treatment by antibiotics and probiotic yeasts: a review. Microbiology, 164(11), pp.1327–1344.

28. Horstmann, J.C., Laric, A., Boese, A., Yildiz, D., Röhrig, T., Empting, M., Frank, N., Krug, D., Müller, R., Schneider-Daum, N. and de Souza Carvalho-Wodarz, C., 2021. Transferring Microclusters of P. aeruginosa Biofilms to the Air–Liquid Interface of Bronchial Epithelial Cells for Repeated Deposition of Aerosolized Tobramycin. ACS Infectious Diseases, 8(1), pp.137–149.

29. Huttner, A.; Harbarth, S.; Carlet, J.; Cosgrove, S.; Goossens, H.; Holmes, A.; Jarlier, V.; Voss, A.; Pittet, D. Antimicrobial resistance: A global view from the 2013 World Healthcare-Associated Infections Forum. Antimicrob. Resist. Infect. Control 2013, 2, 31. https://doi.org/10.1186/2047-2994-2-31.

30. Hussein, M., Allobawi, R., Levou, I., Blaskovich, M.A., Rao, G.G., Li, J. and Velkov, T., 2022. Mechanisms Underlying Synergistic Killing of Polymyxin B in Combination with Cannabidiol against Acinetobacter baumannii: A Metabolomic Study. Pharmaceutics, 14(4), p.786.

31. Ingledew, W.J. and Poole, R.K., 1984. The respiratory chains of Escherichia coli. Microbiological reviews, 48(3), pp.222–271.

32. J Clin Trials.S14:002. Kim, J., 2006. Roles of intermediate conformation and transient disulfide bonding on native folding of P22 tailspike protein. University of Delaware.

33. Janecki, M., Graczyk, M., Lewandowska, A.A. and Pawlak, Ł., 2022. Anti-Inflammatory and Antiviral Effects of Cannabinoids in Inhibiting and Preventing SARS-CoV-2 Infection. International Journal of Molecular Sciences, 23(8), p.4170.

34. Kicman, A. and Toczek, M., 2020. The effects of cannabidiol, a non-intoxicating compound of cannabis, on the cardiovascular system in health and disease. International journal of molecular sciences, 21(18), p.6740.

35. Kim, J., 2006. Roles of intermediate conformation and transient disulfide bonding on native folding of P22 tailspike protein. University of Delaware.

36. Kosgodage, U.S., Matewele, P., Awamaria, B., Kraev, I., Warde, P., Mastroianni, G., Nunn, A.V., Guy, G.W., Bell, J.D., Inal, J.M. and Lange, S., 2019. Cannabidiol is a novel modulator of bacterial membrane vesicles. Frontiers in cellular and infection microbiology, p.324.

37. Lallai, V., Chen, Y.C., Roybal, M.M., Kotha, E.R., Fowler, J.P., Staben, A., Cortez, A. and Fowler, C.D., 2021. Nicotine e-cigarette vapor inhalation and self-administration in a rodent model: Sex-and nicotine delivery-specific effects on metabolism and behavior. Addiction biology, 26(6), p. e13024.

38. Laxminarayan, R.; Duse, A.; Wattal, C.; Zaidi, A.K.; Wertheim, H.F.; Sumpradit, N.; Vlieghe, E.; Hara, G.L.; Gould, I.M.; Goossens, H.; et al. Antibiotic resistance-the need for global solutions. Lancet Infect. Dis. 2013, 13, 1057–1098.

39. Liu, C., Puopolo, T., Li, H., Cai, A., Seeram, N.P. and Ma, H., 2022. Identification of SARS-CoV-2 Main Protease Inhibitors from a Library of Minor Cannabinoids by Biochemical Inhibition Assay and Surface Plasmon Resonance Characterized Binding Affinity. Molecules, 27(18), p.6127.

40. López, R.; García, E.; García, P. Enzymes for anti-infective therapy: Phage lysins. Drug Discov. Today Ther. Strateg. 2004, 1, 469–474. [Google Scholar] [CrossRef].

41. Marini, E.; Magi, G.; Mingoia, M.; Pugnaloni, A.; Facinelli, B. Antimicrobial and anti-virulence activity of capsaicin against erythromycin-resistant, cell-invasive group A streptococcus. Front. Microbiol. 2015, 6, 1281. [CrossRef].

42. Martinenghi, L.D., Jønsson, R., Lund, T. and Jenssen, H., 2020. Isolation, purification, and antimicrobial characterization of cannabidiolic acid and cannabidiol from Cannabis sativa L. Biomolecules, 10(6), p.900.

43. McGrail, J., Martín-Banderas, L. and Durán-Lobato, M., 2022. Cannabinoids as Emergent Therapy Against COVID-19. Cannabis and Cannabinoid Research.

44. Mlost, J., Bryk, M. and Starowicz, K., 2020. Cannabidiol for pain treatment: focus on pharmacology and mechanism of action. International journal of molecular sciences, 21(22), p.8870.

45. Murray, C.J.; Ikuta, K.S.; Sharara, F.; Swetschinski, L.; Aguilar, G.R.; Gray, A.; Han, C.; Bisignano, C.; Rao, P.; Wool, E.; et al. Antimicrobial Resistance, Global burden of bacterial antimicrobial resistance in 2019: A systematic analysis. Lancet 2022, 399.

46. Nguyen, L.C., Yang, D., Nicolaescu, V., Best, T.J., Gula, H., Saxena, D., Gabbard, J.D., Chen, S.N., Ohtsuki, T., Friesen, J.B. and Drayman, N., 2022. Cannabidiol inhibits SARS-CoV-2 replication through induction of the host ER stress and innate immune responses. Science Advances, 8(8), p. eabi6110.

47. O’Neill, J. Tackling Drug-Resistant Infections Globally: Final Report and Recommendations. Government of the United Kingdom.

48. Peyravian, N., Deo, S., Daunert, S. and Jimenez, J.J., 2022. The Anti-Inflammatory Effects of Cannabidiol (CBD) on Acne. Journal of Inflammation Research, 15, p.2795.

49. Pottenger, L.H., Carney, E.W. and Bartels, M.J., 2001. Dose-dependent nonlinear pharmacokinetics of ethylene glycol metabolites in pregnant (GD 10) and nonpregnant Sprague-Dawley rats following oral administration of ethylene glycol. Toxicological Sciences, 62(1), pp.10–19.

50. Qin, X., Yang, M., Cai, H., Liu, Y., Gorris, L., Aslam, M.Z., Jia, K., Sun, T., Wang, X. and Dong, Q., 2022. Antibiotic Resistance of Salmonella Typhimurium Monophasic Variant 1, 4, [5], 12: i: -in China: A Systematic Review and Meta-Analysis. Antibiotics, 11(4), p.532.

51. Rahman, M.U., Wang, W., Sun, Q., Shah, J.A., Li, C., Sun, Y., Li, Y., Zhang, B., Chen, W. and Wang, S., 2021. Endolysin, a promising solution against antimicrobial resistance. Antibiotics, 10(11), p.1277.

52. Rogers, A.W., Tsolis, R.M. and Bäumler, A.J., 2021. Salmonella versus the microbiome. Microbiology and Molecular Biology Reviews, 85(1), pp. e00027–19.

53. Russell, J.B., 2007. The energy spilling reactions of bacteria and other organisms. Microbial Physiology, 13(1-3), pp.1–11.

54. Seltzer, E.S., Watters, A.K., MacKenzie Jr, D., Granat, L.M. and Zhang, D., 2020. Cannabidiol (CBD) as a promising anti-cancer drug. Cancers, 12(11), p.3203.

55. Silmore, L.H., Willmer, A.R., Capparelli, E.V. and Rosania, G.R., 2021. Food effects on the formulation, dosing, and administration of cannabidiol (CBD) in humans: A systematic review of clinical studies. Pharmacotherapy: The Journal of Human Pharmacology and Drug Therapy, 41(4), pp.405–420.

56. Smith, R.; Coast, J. The true cost of antimicrobial resistance. BMJ 2013, 346, f1493. https://doi.org/10.1136/bmj.f1493.

57. Syroeshkin, A.V., Bakeeva, L.E. and Cherepanov, D.A., 1998. Contraction transitions of F1-F0 ATPase during catalytic turnover. Biochimica et Biophysica Acta (BBA)-Bioenergetics, 1409(2), pp.59–71.

58. Tagliaferri, T.L., Jansen, M. and Horz, H.P., 2019. Fighting pathogenic bacteria on two fronts: phages and antibiotics as combined strategy. Frontiers in cellular and infection microbiology, 9, p.22.

59. Tielens, A.G. and Van Hellemond, J.J., 1998. The electron transport chain in anaerobically functioning eukaryotes. Biochimica et Biophysica Acta (BBA)-Bioenergetics, 1365(1-2), pp.71–78.

60. Torres-Barceló, C. and Hochberg, M.E., 2016. Evolutionary rationale for phages as complements of antibiotics. Trends in microbiology, 24(4), pp.249–256.

61. Wassmann, C.S., Højrup, P. and Klitgaard, J.K., 2020. Cannabidiol is an effective helper compound in combination with bacitracin to kill Gram-positive bacteria. Scientific reports, 10(1), pp.1–12.

62. Wassmann, Claes Søndergaard, Andreas Pryds Rolsted, Mie Cecilie Lyngsie, Sergi Torres-Puig, Tina Kronborg, Martin Vestergaard, Hanne Ingmer, Steen Plesner Pontoppidan, and Janne Kudsk Klitgaard. “The menaquinone pathway is important for susceptibility of Staphylococcus aureus to the antibiotic adjuvant, cannabidiol.” Microbiological Research 257 (2022): 126974.

63. Zhou, S., Cui, Z. and Urban, J., 2011. Dead cell counts during serum cultivation are underestimated by the fluorescent live/dead assay (Vol. 6, No. 5, pp. 513–518). Weinheim: WILEY-VCH Verlag.

